# The Cousa objective: a long working distance air objective for multiphoton imaging *in vivo*

**DOI:** 10.1101/2022.11.06.515343

**Authors:** Che-Hang Yu, Yiyi Yu, Liam M. Adsit, Jeremy T. Chang, Jad Barchini, Andrew H. Moberly, Hadas Benisty, Jinkyung Kim, Brent K. Young, Kathleen Heng, Deano M. Farinella, Austin Leikvoll, Rishaab Pavan, Rachel Vistein, Brandon R. Nanfito, David G. C. Hildebrand, Santiago Otero-Coronel, Alipasha Vaziri, Jeffrey L. Goldberg, Anthony J. Ricci, David Fitzpatrick, Jessica A. Cardin, Michael J. Higley, Gordon B. Smith, Prakash Kara, Kristina J. Nielsen, Ikuko T. Smith, Spencer LaVere Smith

**Affiliations:** Department of Electrical and Computer Engineering, University of California Santa Barbara, Santa Barbara, CA 93106, USA; Department of Molecular, Cellular, and Developmental Biology, University of California Santa Barbara, Santa Barbara, CA 93106, USA; Max Planck Florida Institute for Neuroscience, Jupiter, FL 33458, USA; Department of Neuroscience, Yale University, New Haven, CT 06511, USA; Department of Otolaryngology, Stanford University School of Medicine, Stanford University, Stanford, CA 94305, USA; Spencer Center for Vision Research, Byers Eye Institute, School of Medicine, Stanford University, Palo Alto, CA 94304, USA; Neurosciences Interdepartmental Program, Stanford University, Stanford, CA 94305, USA; Laboratory of Neural Systems, The Rockefeller University, New York, NY 10065, USA; Laboratory of Neurotechnology and Biophysics, The Rockefeller University, New York, NY, USA and Kavli Neural Systems Institute, Rockefeller University, New York, NY 10065, USA; Department of Molecular and Cellular Physiology, Stanford University School of Medicine, Stanford University, Stanford, CA 94305, USA; Department of Neuroscience, University of Minnesota, Minneapolis, MN 55455; Department of Biomedical Engineering, University of Minnesota, Minneapolis, MN 55455; Department of Molecular and Comparative Pathobiology, and Zanvyl Krieger Mind/Brain Institute, Johns Hopkins University, Baltimore, MD 21218, USA; Solomon H. Snyder Department of Neuroscience, and Zanvyl Krieger Mind/Brain Institute, Johns Hopkins University, Baltimore, MD 21218, USA; Department of Psychology and Brain Sciences, University of California Santa Barbara, Santa Barbara, CA 93106, USA; Neuroscience Research Institute, University of California Santa Barbara, Santa Barbara, CA 93106, USA

## Abstract

Multiphoton microscopy can resolve fluorescent structures and dynamics deep in scattering tissue, but applying this technique *in vivo* can be limited by short working distance water-immersion objectives. Here we present an ultra long working distance (20 mm) air objective called the Cousa objective. It is optimized for performance across multiphoton imaging wavelengths, offers a > 4 mm^2^ field-of-view with submicron lateral resolution, and is compatible with commonly used multiphoton imaging systems. We share the full optical prescription, and report performance including *in vivo* 2-photon and 3-photon imaging in a range of species and preparations.

## 1. Introduction

Multiphoton microscopy of *in vivo* neuronal activity has been transformative for neuroscience, but its application can be complicated due to the limitations of conventional multiphoton objectives [1–6). Conventional microscope objectives with good multiphoton performance often have short working distances (1-10 mm) and/or require water immersion. The short working distance, coupled with the geometry of the objective tip, can complicate imaging in larger mammals [2, 7], requiring excessive tissue removal and large cranial windows, which can exacerbate immune responses and degrade tissue clarity [8], ultimately limiting the imaging depth and the duration of longitudinal imaging. Exotic applications such as imaging through post-objective optics [9] including prisms [10] or in complex surgical preparations can also be hampered by short working distances. Moreover, water immersion can require awkward reservoirs or the use of gels that can lack appropriate refractive indices and harbor air bubbles that degrade image quality.

To address these issues, we designed an ultra long working distance air immersion objective that is not constrained to conventional engineering trade-offs. It has a working distance of 20 mm, an effective focal length of 20 mm (i.e., a 10x magnification) for a field-of-view (**FOV**) of > 4 mm^2^, a numerical aperture (**NA**) of 0.50, and the optics are designed to minimize aberrations for two-photon imaging across a range of imaging configurations. This objective, referred to as the Cousa, was designed to be compatible with commercial two-photon imaging systems, with standard threading and an entrance pupil of 20 mm. The Cousa objective enables and enhances a range of experiments in various species, using commercial off-the-shelf imaging systems.

## 2. Design

### 2.1. Specifications and constraints

The design specifications of the Cousa objective (**Fig. 1a**) were set and balanced primarily around three factors: *(i)* geometric parameters to facilitate use in animal imaging applications, *(ii)* optimization for two-photon imaging across a large FOV with sub-cellular resolving power, and *(iii)* compatibility with commercial two-photon imaging systems.

**Fig. 1.**
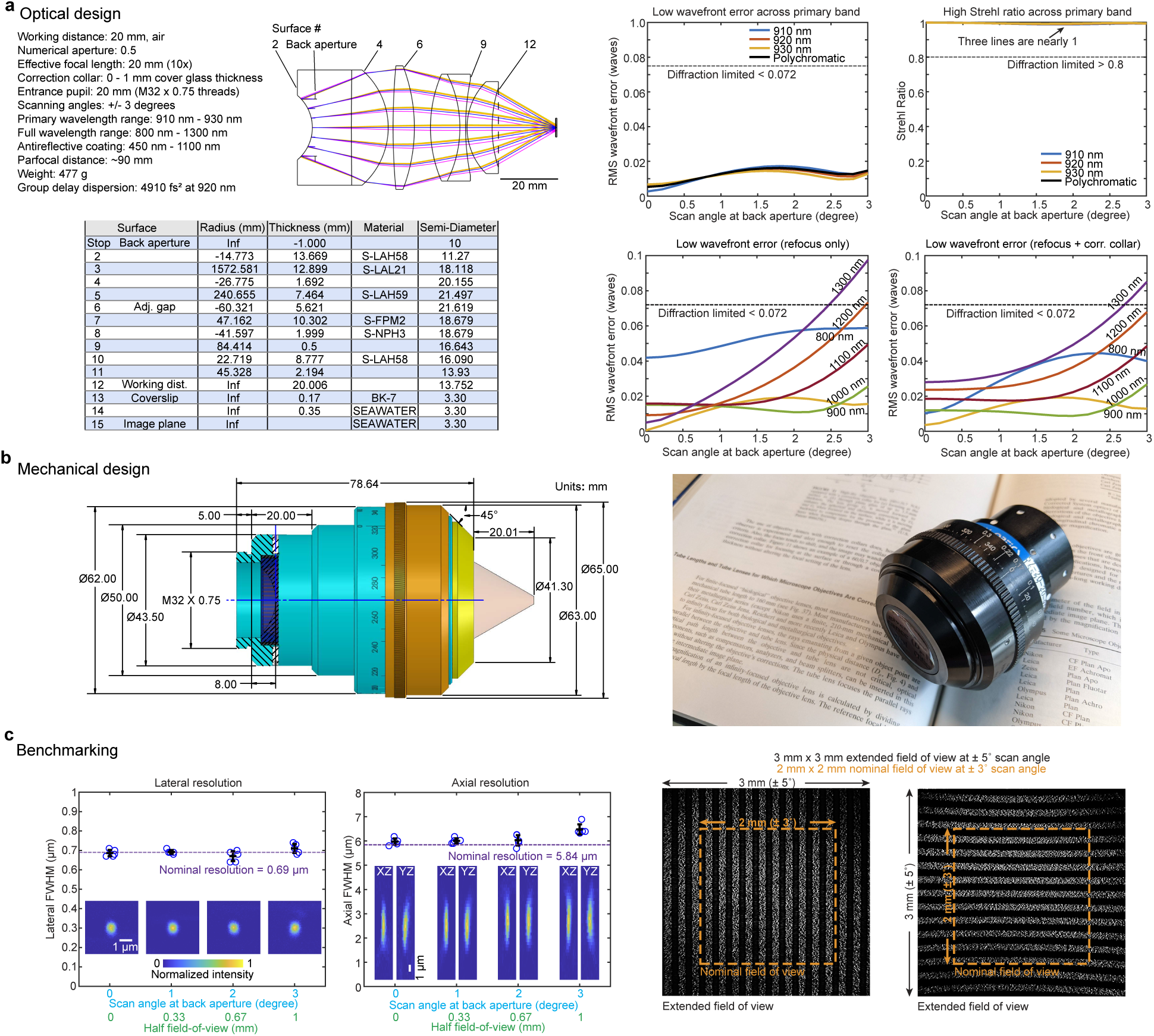
Design and benchmarking. (a, left) The specifications constrained the design to ensure compatibility with two-photon imaging systems that are typically used in vivo. The resulting optical design has six elements and one adjustable air gap to optimize performance. The full lens prescription is provided. (a, right) The primary optimizations were for 920 ± 10 nm, for two-photon excitation of green fluorescent protein (GFP)-based indicators. The optical model predicts low root-mean-squared (RMS) wavefront errors and high Strehl ratios for 910 nm, 920 nm, and 930 nm light across the scan angles of 0 - 3^◦^, well beyond the diffraction limit. Performance is also diffraction-limited across a broader wavelength range from 800 - 1300 nm. The RMS wavefront error remains below the diffraction limit for most of the 0 - 3^◦^ scan angle range, when the focal plane is allowed to naturally shift with wavelength, and the correction collar provides an additional degree of optimization. (b, left) The mechanical design of the objective prioritized keeping the widest diameter near the middle of the objective, to avoid mechanical collisions with objective mounts. All dimensions are in mm unless otherwise noted. (b, right) A photograph of the manufactured objective. (c, left) Two-photon excitation PSF measurements were made with 0.2-*µ*m beads embedded in agar at a depth of 350 *µ*m. The excitation wavelength is 910 nm. Z-stack images are acquired for beads at four lateral locations including on-axis, 1^◦^, 2^◦^, and 3^◦^ off-axis. Full-width at half-maximum (FWHM) of the Gaussian fits for measurements (±S.D.) indicate lateral and axial resolutions indistinguishable from diffraction limited resolutions. The pixel size of the images is 0.058 x 0.064 x 0.69 *µ*m (X x Y x Z). (c,right) Images of a fluorescent calibration sample with a periodic line pattern (5 lines per millimeter) in two orientations acquired under a ±5^◦^ scan angle show a nominal 2 x 2 mm FOV of the objective under the ±3^◦^ scan angle, and a 3 x 3 mm FOV under ±5^◦^ scan angle.

First, a top priority of the design, its *raison d’être*, is compatibility with animal experiments and the associated instrumentation. Two-photon imaging in neuroscience is often performed through a cranial window. The objective is then positioned above the window, at a distance determined by the working distance of the objective. This arrangement can pose constraints on imaging. For example, ferrets and other animals have skulls that are > 1 mm thick with a significant gap between the skull and dura mater. In these cases, cranial windows must be enlarged to accommodate standard two-photon objectives, due to their short working distances and tip geometries. Such large imaging windows create challenges for window positioning, imaging quality, and long-term maintenance. For additional examples, even in smaller animals such as mice, short working distances prevent the insertion of auxiliary optics between the objective and sample, and can also prevent imaging in complex preparations such as ventral access to the cochlea.

To address these issues, we started with the requirement that the working distance would be long, 20 mm. By using a long working distance design, the objective can remain comfortably outside of an imaging chamber, resulting in fewer mechanical constraints. We also recognized that imaging at angles other than the conventional vertical orientation, in particular in larger animals, can make maintaining water immersion difficult. Thus, for this design we chose to use air immersion. Air immersion entails a larger refractive index mismatch than water immersion designs, so we mitigated the trade-off by incorporating a correction collar that can compensate for aberrations (**Fig. 1a**).

Second, the lens design was optimized for focusing ultrafast laser pulses centered at wavelengths commonly used in two-photon imaging, including popular genetically encoded calcium indicators like the GCaMP series [11–13). The optics were designed to offer diffraction-limited performance across a range of wavelengths. We set the NA to be 0.50 (**Fig. 1a**), corresponding to a diffraction-limited resolution of 0.69 *µ*m laterally and 5.84 *µ*m axially, which is sufficient to resolve neurons, dendritic spines, and axonal boutons [14, 15].

Third, the objective was designed to be compatible with commercial multiphoton imaging systems. Major microscope manufacturers use infinity conjugate (i.e., infinity-corrected) optical designs [16], and thus we adopted the same convention for compatibility. The other main constraints from commercial two-photon imaging systems are the beam diameter at the objective back aperture and the maximal scan angle. These parameters are determined by the scan engine. Many commercial two-photon imaging systems constrain the maximal beam diameter to about 20 mm, and the scan angles to about ±3°. We adopted these commonly attained values as design specifications. In addition, we adopted the M32 x 0.75 thread size, which is used on many multiphoton imaging objectives, and thus adapters are readily available (**Fig. 1a**).

With these specifications and constraints set (working distance, air immersion, numerical aperture, scan angle, and back aperture diameter), a relatively large FOV remains feasible by setting the effective focal length of the objective to 20 mm [17]. The design and optimization process was conducted using optical simulation software (ZEMAX OpticStudio). The merit function prioritized maximizing the working distance while maintaining a diffraction-limited point spread function and minimizing the wavefront error across the FOV. Parallel efforts were made to reduce the number of lenses and the thickness of the materials, and thus minimize the size, weight, and cost of the final design.

### 2.2. Design and model performance

The infinity corrected objective consists of six lens elements (**Fig. 1a**) with a net group delay dispersion of ~4910 fs^2^ at 920 nm [18]. The working distance (Surface 12 to the focal plane at Surface 15) is ~20 mm. The position of the back aperture was designed to be very close to the first element, Surface 2. This facilitates alignment in commercial systems, since visual inspection at the back surface can determine whether the excitation beam remains stationary during scanning. Distortions were also minimized across the FOV (**Fig. S1**). Although achromatic performance across visual wavelengths was not a priority (as it commonly is with many conventional widefield and confocal objectives), we also ensured that the collected fluorescence over the visible spectrum of 450-650 nm is focused to a small area (~3 mm) for compatibility with popular photodetectors (**Fig. S2**).

The root-mean-square (**RMS**) wavefront error for 920±10 nm light is less than 0.02*,l* across the scan angles, which is considerably less than the Maréchal criterion of 0.072*,l* (lower is better) (**Fig. 1a**), which is a diffraction limit criterion. Similarly, the Strehl ratio [19] is over 0.97 across the nominal ±3° scan angles (**Fig. 1a**), exceeding the diffraction limit of 0.8 (higher is better). This performance is maintained for at least outside the nominal FOV as well (**Fig. S3**). Thus the performance is diffraction-limited throughout the designed FOV by a clear margin. This margin provides some assurance that performance will remain diffraction-limited despite real-world imperfections that are incorporated during fabrication and assembly.

The objective has an air gap between Surfaces 6 and 7 that is adjusted by a rotating correction collar. Correction collar adjustments can compensate for a range of cover glass (Surface 13) thicknesses, from 0 to 1.0 mm. The correction collar can also be adjusted to optimize performance at different excitation wavelengths (**Fig. 1a**). Adjusting two free parameters, the focal plane location and the correction collar position, diffraction-limited performance can be extended to a range of 800-1300 nm, which covers the commonly-used range of multiphoton excitation wavelengths (**Fig. 1a** and **Fig. S4, S5**). This adjustment can also optimize performance for a range of imaging depths (**Fig. S6a, b**) or samples of varying refractive indices (**Fig. S6c, d**), and coverslips of varying thicknesses (**Fig. S7**). Note that the refocusing is applied at a single position for all scan angles, and the merit function balances performance over the full FOV.

### 2.3. Mechanical model and assembly

After optical designs were finalized, the mechanical design, lens fabrication, housing manufacturing, and objective assembly processes were contracted to an external firm (Special Optics, Denville, NJ, USA). The objective is 79 mm long and 65 mm wide at its widest point. Its total weight is 477 grams (**Fig. 1 a**). The long working distance of the objective relaxes the geometric constraints of the design, as was our strategy. However, one mechanical constraint remained: the objective needed to fit within the clearance around the objective mounting threads of commonly used multiphoton microscopes. A conventional way to load the optics into an objective is to leave the back open, insert all lenses, and then seal it off. This stacking approach leads to the largest diameters being at the back of the objective, near the threads. Realizing this problem in an early version, we redesigned the optomechanics for assembly in the middle, at the adjustable air gap surface (Surface 6). Lenses are loaded from this plane, into both halves, and then the two halves are joined. This reduced the diameter of the shoulder near the threads, and moved the largest diameter to the middle of the lens, where it can be more easily accommodated on commercial multiphoton microscopes. The resulting silhouette of the objective resembles a cousa squash, and inspired the name of the objective (**Fig. 1b**). The total traveling range of the adjustable air gap is 1.0 mm, corresponding to 1.3 revolutions of the correction collar, with a precision of 2.08 *µ*m per degree. The correction collar is marked to indicate both 360-degrees around the objective, and various cover glass thicknesses. The tip of the objective is beveled at 45^◦^ to gain some clearance near the sample space.

## 3. Characterization and performance

### 3.1. Resolution, field-of-view, and light transmission

We characterized the real world performance of the objective using a custom two-photon scan engine with a 32 mm diameter beam scanned over a ±5° range [14]. These scan parameters exceed the requirements of the objective (20 mm and ±3°, respectively), thus the performance should be objective-limited, rather than scan engine-limited [20]. We first measured the resolution attained by the objective by taking z-stacks of 0.2 *µ*m fluorescent beads at various positions across the FOV (**Fig. 1c**). The lateral FWHM is ~0.69 *µ*m throughout the FOV (0.69 ± 0.02 at 0^◦^, 0.69 0.01 at 1^◦^, 0.67 0.03 at 2^◦^, 0.71 0.05 at 3^◦^, mean sd, n=5 at each angle), which is consistent with the theoretical diffraction-limited resolution [21]. The axial resolution is ~5.84 *µ*m, again providing a good match to the theoretical value, up to ±2° scan angles, and deviates by about 10% at 3° of scan angle (5.97 ± 0.13 at 0^◦^, 6.00 0.10 at 1^◦^, 6.03 0.21 at 2^◦^, 6.47 0.21 at 3^◦^, mean sd, n=5 at each angle). The match between the experimental measurement and the theoretical calculation confirms that the NA of the objective is 0.50, as designed (see Methods). This result also implies that the RMS wavefront error is low.

We next measured the imaging FOV with a structured fluorescent sample with periodic lines (5 per mm; item 57–905, Edmund Optics). When the scan angle is ±3°, the images contain 10 lines along both the x and y directions without vignetting, indicating a 2 mm length on each axis of the FOV (**Fig. 1c**). The result demonstrates that the objective has a FOV of 2 × 2 mm^2^ area, consistent with the nominal model performance (**Fig. 1c**). The FOV can be extended to 3 x 3 mm^2^ with a scan of ±5°, and vignetting occurs at the corners of the field (**Fig. 1c**). The distortion is due to a deviation from the F-theta condition in the scanning system of the Diesel2p (**Fig. S1**) [14]. Together, the resolution and FOV provide a space-bandwidth product of (2000 *µ*m / 0.69 *µ*m)^2^ = 8.4 megapixels with ±3° scanning.

### 3.2. Categorical and quantitative comparisons

This objective design is unique in its combination of parameters, which is enabled by being free from the parfocal lengths and mechanical envelopes of conventional objectives. Although the design is compatible with commercial systems, it does not need to conform to the mechanical constraints that standard upright microscopes and confocal objectives must. Free from those design constraints, we could better optimize for the application. For comparison, conventional objectives with NA > 0.3 have a tradeoff between working distance and field-of-view, and the Cousa breaks free from that trade-off (**Fig. 2a**). Similarly, conventional objectives with a FOV > 2 mm^2^ are constrained to working distances < 14 mm. The Cousa breaks free from this trade-off as well (**Fig. 2b**).

**Fig. 2.**
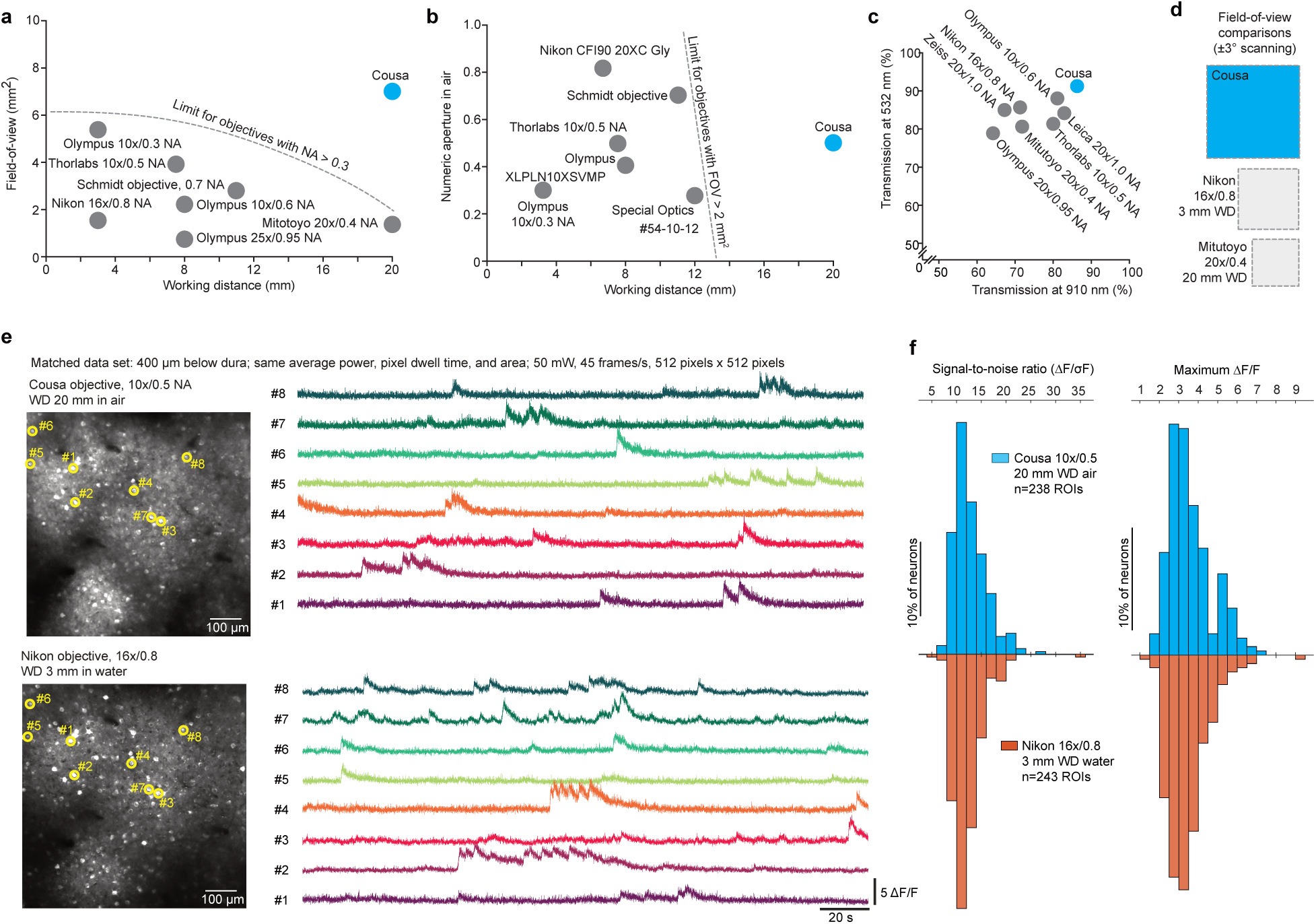
Categorical and quantitative comparisons. (a) Conventional objectives are constrained to a mechanical envelope that limits the product of field-of-view (FOV) and working distance (WD). The general limits imposed by this mechanical envelope are sketched with dotted lines. The Cousa objective is distinctive in that it combines an ultra long WD of 20 mm with a large FOV and an NA of 0.5. (b) Conventional objectives that can offer a FOV > 2 mm^2^ are typically constrained to shorter WD. The Cousa has a long WD, and provides > 4 mm^2^ FOV. (c) Overall throughput in both near IR (910 nm) and visible (532 nm) wavelengths is higher with the Cousa than conventional objectives. This is potentially due to a lower number of lenses in the Cousa. (d) The FOV of the Cousa dwarfs both a commonly used short WD low magnification multiphoton objective (Nikon 16x/0.8NA) and a conventional long WD objective (Mitutoyo 20x/0.4NA). (e) *In vivo* calcium dynamics are measured with the Cousa objective and a popular water immersion objective (Nikon 16x/0.8NA) and compared. The exact same imaging parameters are used to image the same field of neurons in the same mouse (awake, spontaneous activity). Averaged images for both objectives are shown on the left. Calcium traces from the identical region of interests (ROI) are shown on the right (raw data at 45 frames/s; no filtering). The data quality of the Cousa objective is similar to that of this commonly used short WD objective (Nikon 16x/0.8NA). (f) The signal-to-noise ratio (SNR) and maximum f1F/F of the calcium dynamic for each ROI in (e) are calculated and plotted as histograms for both objectives. The signal-to-noise ratio for each ROI is the ratio of the maximal magnitude of the calcium trace to the standard deviation of the fluctuating signal around the baseline. The maximum f1F/F for each ROI is the maximal value of f1F/F throughout the trace. Thus, the Cousa provides an ultra long WD in air, a large field-of-view, and provides raw data similar to that of conventional objectives.

The broadband antireflective coating applied to the lenses was measured to transmit on average 99.5% of visible and near-infrared light (450 - 1100 nm) per surface. To measure the total transmission of 910 nm and 532 nm light through this objective, we supplied an under-filling laser beam into the objective, and measured its power before and after the objective. We found that 86% of 910 nm and 91% of 532 nm light were transmitted through the objective, showing very high transmission throughput at both the excitation and emission wavelengths for green fluorescence protein imaging, especially compared to other commonly used multiphoton imaging objectives (**Fig. 2c**).

The FOV of the Cousa extends past its nominal 2 mm x 2 mm at 3deg scanning. However, we can use this specification for comparison. At those same scan angles, a commonly used short working distance water immersion objective (Nikon 16x/0.8NA) provides smaller FOV. A conventional objective that offers a 20 mm WD in air (Mitutoyo 20x/0.4NA) is limited to an even smaller FOV (**Fig. 2d**, **Fig. S8**).

While the 20 mm WD and large FOV are categorically new to neuroscience, we sought to quantify any decrement in signal quality that might come from the use of the Cousa. We compared performance to a popular short working distance water immersion objective with a higher NA (Nikon 16x/0.8NA). When using the same average power (50 mW), the same pixel dwell times, FOV, and pixel counts, the calcium signals measured *in vivo* at a depth of 400 *µ*m were difficult to distinguish between the two objectives (**Fig. 2e,f**), thus providing confidence that high fidelity data could be obtained with the Cousa. The similarity of the data between the two objectives was likely due to several parameters. On the excitation side, the Nikon 16x/0.8NA objective probably did not achieve the full NA due to the loss of marginal rays in the tissue, while the lower NA Cousa may have experienced less loss of NA [22, 23]. On the detection side, the longer focal length of the Cousa, and thus a larger FOV to collect scattered (non-ballistic) photons, may have partially offset its lower NA. Although the same average power was used out of the objective, on the collection side, the slightly higher transmission of the Cousa would also help make up the difference. Overall, the Cousa provides high fidelity data *in vivo*, 400 *µ*m deep into densely labeled tissue, that compares well with a popular short working distance, higher NA objective (**Fig. 2e,f**).

### 3.3. In vivo two-photon calcium imaging of mouse neural circuitry

After benchmarking and validating the optical performance of the objective, the Cousa objective was used on a range of multiphoton imaging systems, including custom-built systems and commercial systems from Bruker, Thorlabs, Neurolabware, and Sutter. These systems vary in their scan engine performance (e.g., beam size can vary and affect resolution [20]), and thus can provide examples of realistic performance of the objective in experiments. As a first test, a cranial window was implanted in a transgenic mouse with neurons expressing the genetically encoded calcium indicator GCaMP6s [24]. The Cousa objective was mounted on a custom microscope that provided a ±2.6° scan angle range and a 20 mm diameter beam at the back aperture of the objective. This system also had a 12 kHz resonant scanner for fast raster scanning. First, a z-stack image series was acquired covering the volume of 1 x 1 x 0.5 mm^3^ (X*Y*Z) (**Video 1**). This data demonstrated that individual neurons were resolved up to the depth of 0.5 mm. Next, a FOV of 1.7 x 1.7 mm^2^ was recorded using the full ±2.6° scan angle at back aperture (**Fig. 3a**, **Video 2**) with 1536 scan lines, 1536 pixels per scan line, and a frame rate of 15.4 frames/s. Spontaneous calcium transients at a imaging depth of 250 *µ*m were imaged from 1648 neurons detected throughout the FOV (**Fig. 3a**). Calcium indicator traces from neurons across the FOV exhibited high ΔF/F signals (**Fig. 3a**). These *in vivo* results demonstrate performance in the target application, with a relatively large FOV, even when using relatively short pixel dwell times (~28 ns / pixel or ~2 pulses / pixel).

**Fig. 3.**
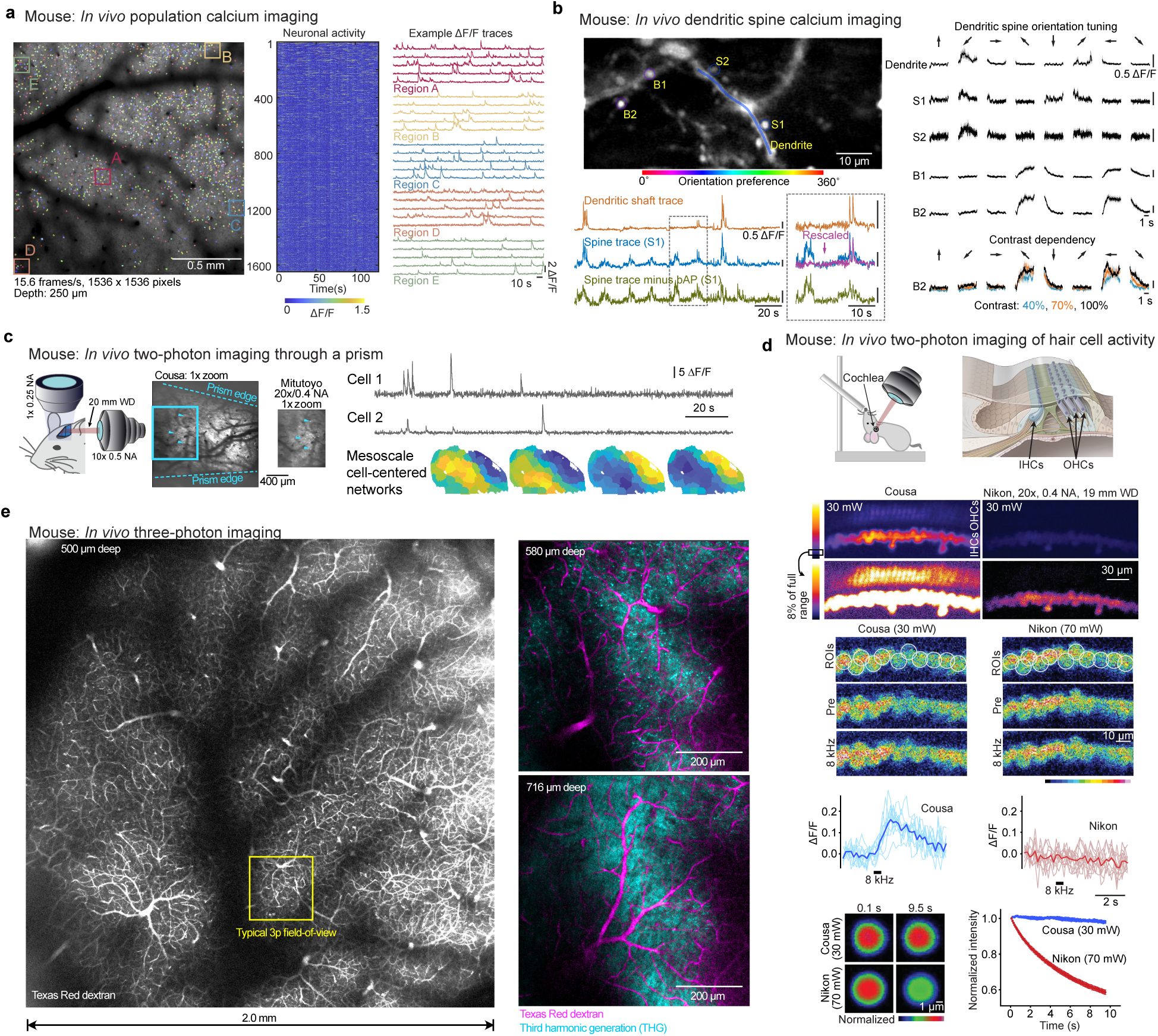
Two-photon and three-photon imaging in mice. (a) Population calcium imaging (GCaMP6s) over a 1.7 mm wide FOV. Traces from cells within boxes at left are expanded at right, a selection of the 1648 neurons detected in this dataset. (b) In a mouse with ultra-sparse expression of GCaMP8m in V1, we imaged calcium transients in putative axonal boutons (B), dendritic spines (S), and dendritic shafts during the presentation of visual stimuli (drifting gratings). Color-codes show the orientation preference of each region of interest. Backpropagating action potential (bAP)-associated calcium transients detected in the dendrite were subtracted from the dendritic spine S1 signal, revealing activity events that are independent from local bAP signals, indicative of local synaptic input. Orientation tuned responses were reliable for spines S1 and S2, boutons B1 and B2, and the nearby dendrite (n = 15 repeats per stimulus; mean in black ± SEM in gray). Responses in axonal bouton B2 varied with contrast (contrast levels of 40% in blue, 70% in orange, and 100% in black (mean ± SEM; n = 5 repeats) (c) Simultaneous two-photon imaging (through a prism) and mesoscopic widefield imaging in awake, head-fixed mice obtained a larger FOV with the Cousa. Time-averaged 2-photon image obtained through the prism show the difference in field-of-view compared to a commercially available 20 mm air objective. Time series for neurons imaged with two-photon excitation through the Cousa objective and microprism were used to detect cell-centered networks for 74 neurons in one mouse. (d, top) The Cousa objective enables *in vivo* functional imaging of cochlear hair cells. The mouse is held supine for imaging of inner hair cells (IHCs) and outer hair cells (OHCs). We imaged with both the Cousa objective and a conventional objective, the Nikon 20x/0.4NA, 19 mm WD (TU Plan ELWD 20X), with the same laser power (30 mW). (d, middle) For calcium imaging in IHCs, the Cousa was used with 30 mW and the Nikon was used with 70 mW, to obtain minimal usable signal levels for both. In response to sound stimulation, IHCs exhibited responses only during Cousa imaging. (d, bottom) Fluorescent particles (diameter 5.63 µm) faded rapidly when imaged with the Nikon, and maintained fluorescence when imaged with the Cousa. (e, left) The Cousa supports large field-of-view 3-photon imaging, with a 20 mm WD. The vasculature across the entire 2 × 2 mm region is visible after an intravenous injection of Texas Red dextran. (e, right) Higher zoom single z plane three-photon images from a second mouse with dual channel imaging of Texas red dextran (magenta) and third-harmonic generation (THG, cyan) in cortex and white matter.

We next performed two-photon imaging of dendrites and axons in a mouse that sparsely expressed GCaMP8m. Neuronal activity in the primary visual cortex (**V1**) was imaged while the animal viewed black and white drifting gratings of eight different orientations (0-315^◦^ and 45^◦^ steps). The spines and their local dendritic shaft are clearly resolved, and some putative boutons are identified with distinctive calcium activity (**Fig. 3b**). Dendritic spine transients showed clear independent calcium dynamics in addition to those associated with back-propagating action potentials (**bAP**), demonstrating that the fluorescence signals from the spine and its parent dendrites can be unambiguously extracted such that the bAP signals can be removed from the spine with high fidelity (**Fig. 3b**) [11]. These identified spines, boutons, and dendrite show reliable response to visual stimuli and different orientation tuning (**Fig. 3b**). Moreover, response magnitude of the axonal bouton transients showed contrast-dependence (40%, 70%, and 100%), further highlighting the sensitivity of the objective and performance in challenging experiments (**Fig. 3b**). Taken together, these results demonstrate that the Cousa objective has not only high resolution for resolving minute structures in neural processes, but also sufficient two-photon excitation and collection efficiency to detect fine changes in calcium transients.

Short working distance objectives preclude the implementation of intermediate optics between the objective and the sample, such as prisms, mirrors, and gradient refractive index (GRIN) lenses [9, 10, 25–27]. Long working distance air objectives can enable experiments such as simultaneous mesoscopic and two-photon imaging of neuronal activity using a prism in the post-objective space [10]. In this method, dual asymmetric imaging pathways are used to record the activity of individual neurons relative to ongoing, large-scale dynamics across the dorsal neocortex. The Cousa objective was mounted horizontally and used in conjunction with a micro prism implanted on the cortical surface (**Fig. 3c**) [10]. Compared to previously used instrumentation (Mitutoyo 20x/0.4NA), the Cousa objective offered a larger FOV, enabling the simultaneous imaging of a larger population of neurons for correlation analysis.

### 3.4. In vivo calcium imaging of the mouse cochlea

The peripheral hearing organ, the cochlea, is difficult to reach surgically and optically. Moreover, it is a mechanosensitive, fluid-filled structure, which further complicates surgical preparations and functional imaging [28, 29]. A long working distance objective is required to keep the neighboring hearing structures intact (**Fig. 3d**). Furthermore, an air objective is needed to preserve sound transference through the air-filled middle ear cavity *in vivo*. Water or oil coupling for the objective interrupts effective sound transfer. Here, we directly compare the performance of the Cousa objective with a long working distance Nikon objective (TU Plan ELWD 20X, 0.4NA, 19 mm WD) in this application, formerly a leading objective for this preparation [28]. *In vivo* two-photon cochlear images were first collected with the same laser power (30 mW) in a genetically modified mouse expressing tdTomato in hair cells (HCs). The signal intensity from inner and outer hair cells (IHCs and OHCs) when using the Cousa objective is higher than that from the Nikon objective (**Fig. 3d**). Next, a mouse expressing GCaMP6s selectively in HCs was used to functionally monitor sound-evoked Ca2+ responses in the cochlea *in vivo* (**Fig. 3d**). An 8 kHz pure tone was played for 500 msec to stimulate the IHCs at the imaging window location. While 30 mW laser power was enough to resolve the Ca2+ sensor in IHCs with the Cousa objective, 70 mW power was required for imaging the cells with the Nikon objective. The Cousa objective revealed IHCs responding to the 8 kHz sound stimulation (**Fig. 3d**). However, we were unable to observe any responsive cells with the Nikon objective (**Fig. 3d**). With the Nikon objective, signals gradually diminished during recording, likely due to the higher laser power used. To examine that possibility, bleaching was induced by the laser powers required for imaging with the Cousa or the Nikon objectives (**Fig. 3d**). The intensity of fluorescent particles decreased rapidly with the Nikon objective and 70 mW power, but were stable with the Cousa objective and 30 mW of power. Taken together, the Cousa objective enabled the first calcium imaging *in vivo* of hair cells, thanks to its unique combination of long working distance and efficiency in two-photon calcium imaging.

### 3.5. In vivo three-photon imaging across a large field-of-view

Next we used the Cousa objective for three-photon (**3p**) imaging. The Cousa supported 3p imaging across the largest field-of-view to date, to the best of our knowledge. We performed three-photon imaging of blood vessels, apical dendrites and white matter axons that were located between 500 – 1,000 *µ*m from the pial surface in mouse visual cortex (**Fig. 3e**, **Supplementary Video 3**). Vessels were labeled with Texas Red dextran, apical dendrites and white matter axons were discernible with label-free third harmonic generation (**THG**) [30–32). Even fine caliber blood vessels such as capillaries were clearly visible in cortical layers 5/6 and within the white matter (**Fig. 3e**). Moreover, in layer 5/6 apical dendrites were clearly represented as bright puncta in the THG channel (**Fig. 3e**, **S9, Supplementary Video 3**). In the white matter, parallel bands of axonal fibers were visible with THG (**Fig. 3e**, **S9, Supplementary Video 3**). These fine-scale structural features located deep in the tissue (puncta of apical dendrites, orientated axonal fibers, capillaries) are reminiscent of the detection of these features when using traditional high NA short working distance water-dipping objective lenses for three-photon imaging, e.g., the Olympus XLPLN25XWMP2, NA 1.05 WD 2 mm (Refs. [30–32)), whose maximum field-of-view is ∼ 750 *µ*m × 750 *µ*m versus ∼ 2,000 *µ*m × 2000 *µ*m with the Cousa.

### 3.6. In vivo two-photon calcium imaging in marmoset

*In vivo* two-photon microscopy in awake monkeys can be particularly challenging. The implants and optical windows are bulky and can physically limit access to the microscope objective, thus a long working distance can be beneficial. Additionally, monkeys can perform tasks more reliably for longer periods of time when they are comfortably upright. In this position, the microscope objective must be positioned more horizontally for accessing many brain areas, and this complicates the maintenance of a liquid immersion interface. Thus, an air immersion objective can facilitate or enable experiments (**Fig. 4a**). We tested the Cousa objective in a marmoset monkey. Two-photon imaging through the Cousa objective was used to resolve neuronal activity of individual neurons expressing jGCaMP7s [12] (**Fig. 4a**). In these marmoset imaging experiments, the setup time was shorter than for water immersion objectives, resulting in more time spent on imaging. The imaging was also more consistent even at extreme cranial window angles because the air immersion eliminated disruptions caused by unstable water or ultrasound gel interfaces or by air bubbles. These experiments show that the Cousa objective enhances multiphoton imaging experiments in awake marmosets.

**Fig. 4.**
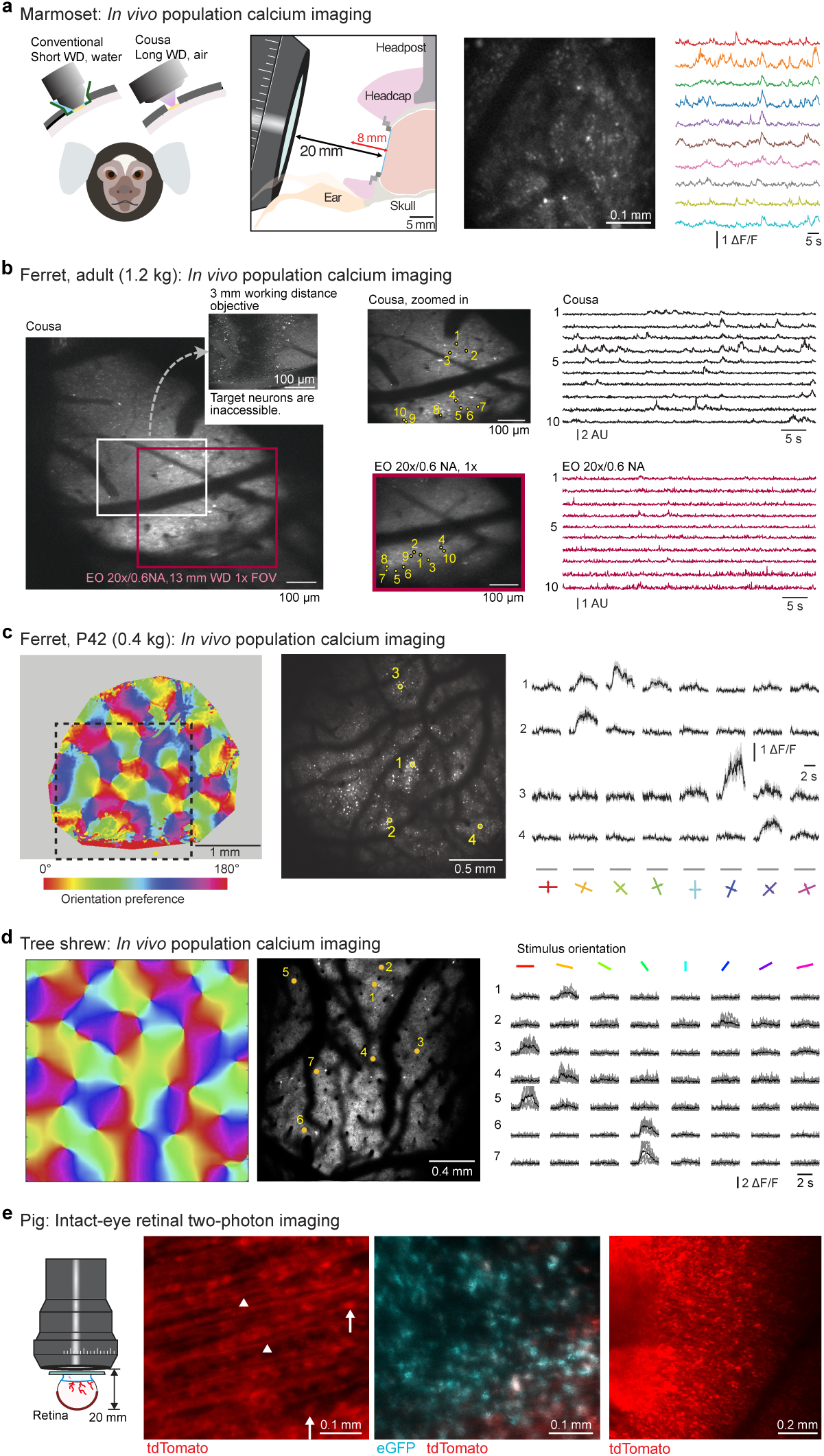
Two-photon imaging in larger mammals. (a) In behaving marmosets, the Cousa objective facilitates imaging replacing water immersion with air and providing ample open space around the cranial window. (b) In ferrets, it can be challenging to access the neurons of interest. A short working distance (WD) objective (Nikon 16x/0.8NA, 3 mm WD in water) cannot even access the neurons of interest due to the short WD and collisions with the walls of the cranial window. The Cousa provides a large FOV and activity measurement, another long working distance air objective provides only a smaller FOV (at a different angle) and weaker signals. (c) In ferrets, with another type of imaging window, the Cousa provides a large field-of-view for measuring orientation tuning across orientation columns. (d) Similarly, in tree shrew, the Cousa provides a large field of view for resolving individual neurons across orientation domains. (e) In a challenging preparation, the Cousa enabled two-photon imaging through the lens and entire eye to the retina of an intact porcine eye, including retinal ganglion cell bodies (arrows) and single axon fibers (triangles).

### 3.7. In vivo two-photon calcium imaging in ferret and tree shrew

Ferrets and tree shrews are important model systems in neuroscience, and these preparations can pose challenges for short working distance water immersion objectives. Thus, we tested the Cousa objective for imaging in these animals. Two labs performed experiments in ferrets. In adult ferrets, the skull is thick and has prominent crests, or ridges, which result in cranial windows with significant geometric constraints. The use of a short working distance objective is impaired by mechanical collisions with the walls surrounding the window. An objective with a 13 mm working distance provided some optical access to the preparation. However, it was still not complete, and the imaging angle had to be altered to accommodate the objective. The imaging quality was relatively poor, especially compared to the Cousa objective, which provided a larger field-of-view, and better data quality (**Fig. 4b**).

For imaging in younger ferrets and tree shrew V1, the objective was mounted on a commercially available microscope (B-Scope; Thorlabs). Injected viral particles transduced neurons to express GCaMP6s [24]. For ferret imaging, wide-field imaging was used to image the vasculature and the orientation preference map in V1 (**Fig. 4c**). Then, two-photon imaging through the Cousa objective was used to resolve neuronal activity of individual neurons within a 2 mm x 2 mm FOV (**Fig. 4c**). Individual neurons exhibited reliable responses to visual stimuli with edges of particular orientations (**Fig. 4c**). Observed two-photon orientation preferences were also consistent with their location within the orientation preference map (as measured with widefield imaging). In tree shrews, V1 neurons were transduced to express GCaMP6s [24]. The Cousa objective was used to image calcium transients (**Fig. 4d**). Individual neurons could be registered to their location in the local orientation preference map (**Fig. 4d**), and reliable responses to visual stimuli were resolved (**Fig. 4d**).

In both ferret and tree shrew V1 imaging, the Cousa objective offered a larger FOV than conventional objectives, and the air immersion facilitated imaging, especially at angles where maintenance of a water interface can be unreliable. Together, these experiments demonstrate that the Cousa objective supports multiphoton imaging in ferrets and tree shrews.

### 3.8. Imaging the porcine eye

In studies of the neural circuitry of the porcine eye, cells are virally manipulated to express fluorescent indicators. However, imaging these labeled cells with single-cell resolution typically requires the excision of the retina, which disrupts circuitry and precludes longitudinal studies. We used the Cousa objective to image cells and axons in the intact porcine eye, despite its ~20 mm diameter. First, we injected AAV particles to express eGFP and tdTomato in porcine retina *in vivo*. Next, the eye was placed under the objective with hydroxypropyl methylcellulose gel (Ocular Vision) and coverslipped to match the corneal refractive index and flatten this portion of the optical pathway (**Fig. 4e**).

We imaged retinal ganglion cell (RGC) axonal fibers that have a diameter of ∼1 µm and cell somas (**Fig. 4e**). Next, we identified an area of high eGFP and td-Tomato co-localization of RGC somas (**Fig. 4e**), demonstrating the ability of the objective and microscope system to image green and red fluorophores simultaneously. We then imaged an area of ~1 mm^2^ to visualize blood vessel patterns for registering with post-dissection images (**Fig. 4e**). The Cousa objective enabled direct imaging of the porcine retina through the intact eye, resolving individual cells and axon fibers.

## 4. Discussion

In summary, we developed a microscope objective optimized to enable and enhance certain challenging multiphoton imaging experiments. The key attribute of the Cousa is its 20 mm working distance in air. It is unique in that it combines this long working distance with an NA of 0.50, optimization for multiphoton imaging, and a long effective focal length, which provides a FOV of 4 mm^2^ (up to 9 mm^2^ at ± 5^◦^ scanning). The manufactured objective has been distributed to an array of labs, and their results demonstrate functional and structural two-photon, three-photon, and THG imaging *in vivo*. While conventional objectives remain an excellent choice for conventional experiments, there are several noteworthy results we presented here, that were enabled or enhanced by the Cousa objective. We presented the first *in vivo* functional imaging of cochlear hair cells. We presented the first multiphoton imaging of porcine retina, through the intact eye. Finally, we also presented the widest field-of-view 3-photon and THG imaging *in vivo*.

Prior development of an objective like the Cousa was probably impaired by multiple issues. First, conventional microscopes have a mechanical envelope in which the microscope objective must fit. Objectives must fit on turrets and not collide with microscope stages [16]. In this work, we recognized that the field of multiphoton imaging *in vivo* has evolved to support a range of imaging systems, many with no turrets or substage optics, that can accommodate objectives with mechanically larger form factors. Second, conventional microscopy has prioritized broad chromatic corrections and high numerical apertures, and placed a lower priority on working distance and balanced performance across a large field-of-view [16, 33]. Thus, the area of the parameter space where the Cousa lies is relatively underexplored. Third, the potential applications for an ultra long working distance multiphoton objective were unclear. In this study, we show an array of applications that are enabled or enhanced by using a multiphoton-optimized ultra-long working distance objective, including complex surgical preparations and applications with auxiliary optics in the post-objective space, including prisms and gradient index (GRIN) lenses. The Cousa can also enable experiments that entail multiphoton imaging simultaneous with electrophysiology using electrode arrays [34] or patch clamp electrodes [35, 36] where the electrodes would prevent the use of short working distance objectives.

The lens description is open-source, so that the community can replicate, modify, or simulate for their applications. It could be promising to explore various engineering tradeoffs, including working distance, mechanical size, NA, and chromatic corrections. The objective we present here demonstrates the broad utility and impact of alternative optical designs that depart from conventional parameters and constraints.

## 5. Methods

### 5.1. Objective design and assembly

The objective was modeled and optimized using an optical design software of OpticStudio (Zemax, LLC). Tolerance analysis indicated that 90% of the completed objectives would have an RMS wavefront error of 0.048*λ* (still well below the diffraction limit criterion of 0.072*λ*) with commonly attained manufacturing and assembly tolerances. All lenses in the objective were manufactured, aligned, and assembled in the factory of Special Optics (Denville, NJ, USA). The manufacturing tolerances used were 0.005 mm total indicator runout (TIR) for decentration and tilt, 0.05 mm for thickness, 4 rings for radius (power), .25 waves at 633 nm for irregularity, 0.005 mm for wedge, 60-40 scratch-dig, and 0.01 arc min for lens decentration.

### 5.2. Custom in vivo two photon imaging system

Two custom two-photon systems were used for most of the characterization and the mouse experiments in Fig. 1, 2, and 3a,b. One system was equipped with an 8 kHz resonant scanner (CRS 8 kHz, Cambridge technology) and dual galvo scanners and supplied a 32-mm diameter beam size and 5^◦^ scan angles at the objective back aperture [14]. The other system was equipped with a 12 kHz resonant scanner (CRS 12 kHz, Cambridge technology) and supplied a higher imaging frame rate with a 20-mm beam size and 2.6^◦^ scan angles at the objective back aperture. Our laser source was a Ti:sapphire pulsed laser with a central wavelength at 910 nm and an 80 MHz repetition rate (Mai-Tai, Newport). The image acquisition was controlled by ScanImage (Vidrio Technologies). The imaging was performed with a power ≤80 mW out of the front of the objective. Other imaging systems were used for the data in Fig. 3 and 4 and are detailed in *Section 5.4* below.

### 5.3. Excitation point spread function measurements and simulations

The measurement and analysis procedure were described in our previous publication in details [15]. To evaluate the excitation point spread function (PSF), sub-micrometer beads were imaged. Sub-micrometer fluorescent beads (0.2 *µ*m, Invitrogen F-8811) were embedded in a thick (~1.2 mm) 0.75% agarose gel. 30 *µ*m z-stacks were acquired, each centered at a depth 350 *µ*m. The stage was moved axially in 0.5 *µ*m increments (Δstage). At each focal plane 30 frames were acquired and averaged to yield a high signal-to-noise image. Due to the difference between the refractive index of the objective immersion medium (air) and the specimen medium (water), the actual focal position within the specimen was moved an amount f1focus = 1.38 x f1stage [37]. The factor 1.38 was determined in Zemax and slightly differs from the paraxial approximation of 1.33. These z-stack images were imported into MATLAB for analysis. For the axial PSF, XZ and YZ images were created at the center of a bead, and a line plot was made at an angle maximizing the axial intensity spread, thereby preventing underestimation of the PSF due to tilted focal shifts. For the radial PSF, an XY image was found at the maximum intensity position axially. A line scan in X and Y was made. Gaussian curves were fit to the individual line scans to extract FWHM measurements. The radial PSF values are an average of the X PSF and Y PSF, and the axial PSF is an average of the axial PSF found from the XZ and YZ images. Excitation PSF measurements were performed at locations of on axis, 1^◦^, 2^◦^, and 3^◦^ off axis across the FOV. Data reported (Fig. 1c) are the mean ± S.D. of 5 beads (n=5) at each location.

The theoretical calculation of the PSF is based on the equations as follows, and can be converted into FWHM by multiplying 2 · 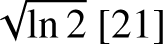.

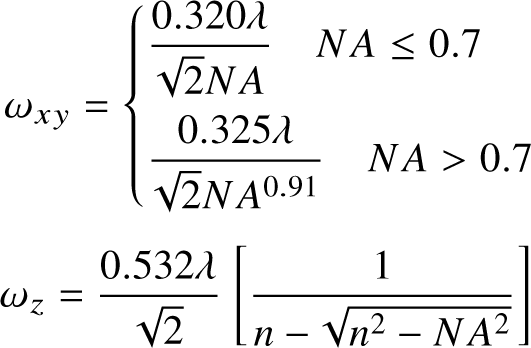

NA is the numerical aperture of the objective lens, n is the refractive index of the medium where the sample is embedded, and *,l* is the excitation wavelength. Using the value of 910 nm for the excitation wavelength, 0.5 for the NA, and 1.33 for the refractive index of water, we have FWHM*_xy_* = 0.69 *µ*m, and FWHM*_z_* = 5.84 *µ*m.

### 5.4. Animal experiments

#### 5.4.1. Mouse: Population calcium imaging and dendritic calcium imaging

All procedures involving living animals for these figures were carried out in accordance with the guidelines and regulations of the US Department of Health and Human Services and approved by the Institutional Animal Care and Use Committee at University of California, Santa Barbara. Mice were housed in 12 h dark/light reverse cycle room. The temperature set-point is 74–76 ^◦^F; the low-temperature alarm is 70 ^◦^F; the high-temperature alarm is 78 ^◦^F. The relative humidity is 45% (range 30–70%).

For population calcium imaging, GCaMP6s transgenic mice were used as before [38], which were generated by triple crossing of TITL-GCaMP6s mice, Emx1-Cre mice (Jackson Labs stock #005628) and ROSA:LNL:tTA mice (Jackson Labs stock #011008). TITL-GCaMP6s mice were kindly provided by Allen institute. Transgenic mice were deeply anesthetized using isoflurane (1.5–2%) augmented with acepromazine (2 mg/kg body weight) during craniotomy surgery.

Carpofen (5 mg/kg body weight) was administered prior to surgery, as well as after surgery for 3 consecutive days. Glass windows were implanted over visual cortex as previously described [14, 38]. Ca^2+^ signals were analyzed using custom software [39] in MATLAB (Mathworks). Neurons were segmented and fluorescence time courses were extracted from imaging stacks using Suite2p (https://suite2p.readthedocs.io/en/latest/) [40]. Signals from neurons are a sum of neuronal and neuropil components. The neuropil component was subtracted from the neuronal signals by separately detecting it and subtracting it. The neuropil component was isolated using the signal from an annulus region around each neuron, and then subtracted from the neuronal signal to provide a higher fidelity report of neuronal fluorescence dynamics. An exponential moving average with a moving window size of 5 samples (0.32 s) was used to reduce the baseline noise in the traces displayed (**Fig. 3a**).

For dendrite calcium imaging, adult (> 8 weeks) C57Bl/6 mice of both sexes (Jackson Labs) were used. A 4-mm diameter craniotomy was performed over visual cortex as previously described [41]. Briefly, mice were premedicated with a sedative, acepromazine (2 mg/kg body weight, i.p.), after which they were deeply anesthetized using isoflurane (2-3% for induction, 1-1.5% for surgery). The mouse’s body temperature was monitored and actively maintained using an electronic heat pad regulated via rectal probe. Carprofen (5 mg/kg body weight, s.c.) was administered preoperatively, and lidocaine solution containing epinephrine (5 mg/kg body weight s.c.) was injected locally before and after the scalp excision. The scalp overlaying the right visual cortex was removed and a custom head-fixing imaging chamber with a 5-mm diameter opening was mounted to the skull with cyanoacrylate-based glue (Oasis Medical) and dental acrylic (Lang Dental). Mice were mounted on a custom holder via the headplate chamber, which was filled with a physiological saline containing (in mM) 150 NaCl, 2.5 KCl, 10 HEPES, 2 CaCl2 and 1 MgCl2. A craniotomy was performed using carbide and diamond dental burs on a contra-angle handpiece (NSK). adeno-associated viral (AAV) vectors were injected in to V1 under continued isoflurane anesthesia as previously described [41–43). Briefly, 1:1 mixture of pENN.AAV.CamKII 0.4.Cre.SV40(AAV1; Addgene #105558; diluted at 1:20,000 in phosphate buffered saline (PBS)) and pGP.AAV.syn.FLEX.jGCaMP8m.WPRE (AAV1; Addgene #162378; original concentration at 10^13^ vg/mL) viral particles were injected (80 nL per site; 1 site per animal) into V1 with a pulled-glass capillary micropipette using a Nanoliter 2010 controlled by a microprocessor, Micro4 (World Precision Instruments), at 15 nL per min. The glass pipette was left in place for 5 mins before retracting to avoid the backflushing of the injected solution. The cranial window was then sealed with a glass cranial plug made up of 4-mm and 3-mm circular coverslips (Warner Instruments) stacked in tandem with a UV-curing optical adhesive (NOA61, Norland). Two-photon imaging of Ca2+ transients indicated by GCaMP8m was performed starting 4-6 weeks after AAV injection, using a custom-built two-photon microscope used in prior studies [35, 41]. Frame scans were acquired using ScanImage [44] at 58.2 frames per second, 512×256 pixels; 31000 frames total per visual stimulation session.

Visual stimuli were presented on a 7” monitor (60 Hz refresh rate) placed 12 cm away from the animal’s eye. To assess orientation tuning of the dendritic shaft, spines, and putative axonal boutons, full field square gratings at 40%, 70%, and 100% contrasts (0.04 cycles per degree at 2 Hz) were presented in 8 directions (0^◦^, 45^◦^, 90^◦^, 135^◦^, 180^◦^, 225^◦^, 270^◦^, 315^◦^) for 5 trials. Each grating drifted for 4 seconds. A notch filter centered at 2 Hz (±0.5 Hz bandwidth) was used to remove a small amount of light leakage from the stimulus monitor into the imaging pathway.

To functionally map visual cortex for targeted injection of viral vectors, ISOI was performed using a custom macroscope and a CCD camera as previously described [41, 45]. Retinotopic maps were used to locate V1. The pial vasculature map relative to the retinotopic maps was used to guide targeted injections into V1.

#### 5.4.2. Mouse: Cochlea imaging

Animal studies were carried out according to the protocols approved by the Institutional Animal Care and Use Committee at Stanford University (APLAC-14345). Four weeks old male mouse from Ai14tdTomato (JAX: 007908) x Myosin15Cre [46] breeding was used for cochlear hair cell imaging. The mouse was anesthetized using ketamine (100 mg/kg) and xylazine (10 mg/kg). Anesthesia level was assessed by signs of movement or withdrawal reflex before the application of supplementary anesthetic. Mouse surgery and positioning for *in vivo* cochlear imaging was performed by the method described previously [29]. *In vivo* cochlear imaging was performed using a modified commercial two-photon microscope (Ultima, Bruker) with long working distance air objectives (Cousa objective; TU Plan ELWD 20X, NA 0.4, WD 19mm, Nikon Instruments Inc.). A Ti:sapphire laser was used with wavelength 920 nm and power 30 mW (Chameleon, Coherent Inc.). The projected images (**Fig. 3d**) were acquired in an apical hair cell location (8-10 kHz) by collecting z-series 40 images with 2 *µ*m intervals.

#### 5.4.3. Mouse: Prism-based two-photon imaging with simultaneous wide-field imaging

All animal handling and experiments were performed according to the ethical guidelines of the Institutional Animal Care and Use Committee of the Yale University School of Medicine. Brain-wide expression of GCaMP6s was achieved via neonatal sinus injection of AAV9-Syn-GCaMP6s into c57/Bl6 mice, as described previously [10, 47]. After reaching adulthood (P60), the skin and fascia over the skull were removed under isoflurane anesthesia and the animal was implanted with a custom titanium headpost and a microprism (5mm per side, Tower Optics) placed over the right visual cortex in a small craniotomy, bonded with a thin layer of dental cement (Metabond, Parkell).

Imaging experiments were carried out in awake mice head-fixed over a freely-moving wheel placed under the microscope objective. Widefield calcium imaging was performed using a Zeiss Axiozoom with a PlanNeoFluar objective (1x, 0.25 NA). Epifluorescent excitation was provided by an LED bank (Spectra X Light Engine, Lumencor) strobing 395 nm and 470 nm light, for hemodynamic correction and calcium imaging, respectively [47]. Emitted light was collected via sCMOS camera (Orca-Flash V3, Hamamatsu), with images acquired at 512 x 512 pixel resolution and 10 frames per second. Data were pre-processed for hemodynamic correction and normalized to f1F/F values as previously described 2. Functional parcellation of cortical areas was carried out using local selective spectral clustering (LSSC, [47, 48]) to obtain a time series of fluorescence signal for each parcel.

Two-photon imaging was performed using a resonant-galvo scanning microscope (MOM, Sutter Instruments) coupled to our custom air-coupled, long-working distance objective (10x, 0.5 NA). Excitation was provided by a titanium-sapphire laser (MaiTai, SpectraPhysics) tuned to 920 nm. Light was directed into the brain after being reflected 90^◦^ by the implanted prism. Emitted light was collected by a gallium arsenide-phosphide detector (Hamamatsu) with images acquired at 512 x 512 pixel resolution and 30 frames per second. Data were motion corrected using NoRMCorre [49], and regions of interest (ROIs) corresponding to single cells were manually selected, neuropil-corrected, and normalized to f1F/F values using custom software written in Matlab (Mathworks).

We calculated cell-centered networks (CCNs) to quantify the relationship between activity in single neurons and the large-scale cortical network in the contralateral hemisphere as described previously [10]. Briefly, we evaluated the correlation coefficients between time series related to p mesoscopic parcels and n time series related to cells to obtain C, a p×n matrix. We viewed each column of C as a compact representation of synchrony between the dynamics of each cell and the dynamics of the widefield signal and then clustered these vectors using the kmeans (k=4) function in Matlab. We obtained the centroid map of each cluster as the average correlation coefficients of all cells related to a specific cluster.

We then superimposed each centroid onto the full cortex parcellation to yield the average images in Fig. 3c.

#### 5.4.4. Mouse: Three-photon imaging

All animal procedures were approved by the Institutional Animal Care and Use Committee at the University of Minnesota. Our surgical and imaging methods have been described previously [31, 50, 51]. Briefly, in testing the Cousa objective lens for deep tissue three-photon imaging, two C57BL/6J mice were used. Mice were initially anesthetized with a bolus injection of fentanyl citrate (0.05 mg kg^-1^), midazolam (5 mg kg^-1^), and dexmedetomidine (0.25 mg kg^-1^). A craniectomy (3–4 mm in diameter) was made over the visual cortex. The cranial windows were sealed with agarose (1.5% in artificial cerebrospinal fluid) and a glass coverslip (5 mm diameter, 0.15 mm thickness; Warner #D263). During imaging, continuous intraperitoneal infusion with a lower concentration mixture (fentanyl citrate: 0.002–0.03 mg kg^-1^ hr^-1^, midazolam: 0.2–3.0 mg kg^-1^ hr^-1^, and dexmedetomidine: 0.010–0.15 mg kg^-1^ hr^-1^) was administered using a catheter connected to a syringe pump. Three-photon imaging was performed using a microscope from Bruker coupled to a fixed wavelength laser source from Class 5 Photonics (Hamburg Germany). The excitation wavelength was fixed at 1300 nm using a White Dwarf WD-1300-80-60 femtosecond optical parametric chirped pulse amplifier (OPCPA) pumped by a 0.5 MHz Coherent Monaco-1035-80-60 laser. The Cousa objective lens correction collar was set to 0.15 mm to compensate for the aberration from the coverslip. Three-photon brightness was maximized using a motorized single-prism pulse compressor (BOA-1300, Swamp Optics). To visualize blood vessels, we injected Texas Red dextran retro-orbitally [31]. In addition, we used the label-free Third Harmonic Generation (THG) imaging at 1300 nm excitation to visualize apical dendrites and white matter axons [30, 31].

#### 5.4.5. Marmoset experiments

All experimental procedures were approved by The Rockefeller University Institutional Animal Care and Use Committee and were performed in accordance with guidelines from the U.S. National Institute of Health. One four-year-old marmoset participated in this study.

A headpost and headcap were first implanted, followed by implantation of a 10 mm-diameter cranial window over temporal cortex. The coverglass was 0.17 mm in thickness. Viral transduction was accomplished with microinjection of a solution containing a 1:1 ratio of AAV2/9:Thy1S-tTA (Vigene Biosciences, 6.57 x 10^13^ gc/mL) and AAV2/9:TRE3G-jGCaMP7s-WPRE (Vigene Biosciences, 6.20 x 10^13^ gc/mL) [52]. The solution had a titer of 1×10^13^ vg/mL for each virus and was injected through glass micropipettes (tip diameter 40 – 60 µm, 30^◦^ bevel) in a volume of 500 nL at 0.5 mm and again at 1.0 mm depths. Thirteen locations across the window were injected at a rate of 100 nL/min with a 10 min pause before retracting the micropipette from each site.

The marmoset was placed into a custom-made chair and head fixation established using a custom-designed triangular headpost fit into a custom-machined clamp. Two-photon imaging of jGCaMP7s was performed with a custom-built microscope equipped with a 12 kHz resonant-galvo scanner and driven by a Spectra-Physics Mai-Tai DeepSee laser tuned to 920 nm at a framerate of 45 Hz or with a Thorlabs Multiphoton Mesoscope driven by a Class5 Photonics White Dwarf laser tuned to 960n m at a frame rate of 6.36Hz. Average excitation power at the exit of the objective ranged from 50 to 80 mW. The objective was angled 70 – 90° relative to the body axis of the marmoset. Microscopes were controlled by ScanImage (MBF Bioscience). Images were acquired in the absence of controlled stimuli. Neuron ROI locations and fluorescence time courses were extracted from the resulting image stacks using Suite2p (https://suite2p.readthedocs.io/en/latest/) [24]. Fluorescence traces for each ROI were calculated as the f1*F F* = *F F*_0_ *F*_0_, where the baseline fluorescence (*F*_0_) was approximated as the mean fluorescence across the entire image stack.

#### 5.4.6. Ferret and tree shrew experiments

All experimental procedures were approved by the Max Planck Florida Institute for Neuroscience Institutional Animal Care and Use committee and were performed in accordance with guidelines from the U.S. National Institute of Health. We used one juvenile female ferret from Marshal Farms and one adult male tree shrew for this study.

Viral transduction and terminal imaging in L2/3 of the anesthetized ferret and tree shrew were performed as previously described [53, 54]. Briefly, we expressed GCaMP6s by direct microinjection of AAV2/1-hSyn-GCaMP6s-WPRE-SV40 (Addgene, 100843-AAV1, Titer: 2.5×10^13^ GC/mL) into the visual cortex. Subsequently a cranial window was implanted over visual cortex and imaged. An injection into the visual cortex of the ferret was made at P21, and imaging was performed at P42 (386 g body weight). Imaging in the tree shrew occurred 16 days after viral transduction.

Two-photon imaging of GCaMP6s was performed with a Bergamo II series microscope (Thorlabs) equipped with an 8 kHz resonant-galvo scanner and driven by a Mai-Tai DeepSee laser or Insight DS+ (Spectra-Physicis) at 910 nm or 920 nm respectively. Average excitation power at the exit of the objective ranged from 40 to 60 mW. The microscope was controlled by ScanImage (MBF Bioscience). Images were acquired at 15 Hz (1024 x 1024 pixels in the ferret, 512 x 512 pixels in the tree shrew). Widefield epifluorescence imaging of GCaMP6s in the ferret was achieved with a Zyla 5.5 sCMOS camera (Andor) controlled by *µ*Manager [55] through a 4x air-immersion objective (Olympus, UPlanFL 4x N/0.13NA) and images were acquired at 15 Hz with 4×4 binning to yield 640×540 pixel images.

Visual stimuli were presented on an LCD screen using PsychoPy (v1.85) [56]. The monitor (30 cm x 52 cm, 1920 x 1080 pixels, 120 Hz refresh rate) was placed 25 centimeters in front of the animal. To evoke orientation-specific responses, full field square gratings at 100% contrast were presented in 16 directions (8 orientations) for 10 trials (ferret) or 8 trials (tree shrew). Square gratings were presented to the ferret at 0.06 cycles per degree and 4 Hz and in the tree shrew at 0.4 cycles per degree and 2 Hz. In addition, “blank” stimuli of 0% contrast were also presented. All stimuli were randomly interleaved and presented for 4s followed by 6s of gray screen (ferret) or 2s followed by 3s of gray screen (tree shrew). Timing for visual stimuli and imaging were recorded using Spike2 (v7.11b, CED; Cambridge UK).

Data analysis in the ferret was performed as previously described using custom written scripts in Python and ImageJ [57]. For both widefield and epifluorescence imaging, we corrected brain movement during imaging by maximizing phase correlation to a common reference frame. In widefield epifluorescence imaging, the ROI was drawn manually around regions where robust visually evoked activity was observed. For analysis, all images were spatially downsampled by a factor of 2 to yield 320×270 pixels. Slow drifts in fluorescence intensity were eliminated by calculating the f1*F F* = *F F*_0_ *F*_0_. Baseline fluorescence (*F*_0_) was calculated by applying a rank-order filter to the raw fluorescence trace (10^th^ percentile) with a rolling time window of 60 s. Responses were filtered with a spatial band-pass filter with low-pass cutoff defined as 50 *µ*m and high-pass filter cutoff as 3200 *µ*m. Preferred orientation was computed by taking the vector sum of the median-trial response over the stimulus period for each orientation.

For analysis, ROI were chosen semi-automatically (Cell Magic Wand v1.0) and fluorescence was computed by averaging all pixels within the ROI [42]. The f1F/F for each ROI was computed, and F0 was calculated by applying a rank-order filter to the raw fluorescence (20^th^ percentile) over a rolling time window (60 s). Stimulus-evoked responses were calculated as the average ΔF/F over the entire stimulus period, and orientation preferences were computed by fitting a von Mises distribution to the trial-median response for each stimulus orientation.

Data analysis and motion correction in the tree shrew was performed using custom code written in Matlab (Mathworks) or Java package for running ImageJ within Matlab (Mĳi). For network-level analysis, the fluorescence signal for each pixel was calculated as ΔF/F, where *F*_0_ is the baseline fluorescence signal averaged over a 1 s period immediately before the start of visual stimulus, and F is the fluorescence signal averaged over the period of the stimulus. Responses to the stimulus set were fitted with a Gaussian to determine the preferred orientation and generate a pixel-based orientation preference map. For analysis at the neuronal level, regions of interest (ROIs) corresponding to visually identified neurons were drawn manually using ImageJ. The fluorescence of each ROI was measured by averaging all pixels within the ROI.

#### 5.4.7. Adult ferret experiments with objective comparisons

All procedures adhered to the guidelines of the National Institute of Health and were approved by the Animal Care and Use Committee at Johns Hopkins University. The experiment was performed in an adult male ferret (Mustela putoris furo; 120 days old, 1.2 kg body weight). Virus was injected during an aseptic procedure under isoflurane anesthesia. During this procedure, a craniotomy of approximately 2×2 mm was made over V1. Virus (AAV1.syn.jGCaMP7s.WPRE.SV40 from Addgene, lot v50167, titer 2.7E13 GC/mL) was then injected at 2 sites within the durotomy. At each site, injections were performed at multiple depths for a total of about 1uL per site. 14 days after the procedure to allow for virus expression, we then performed an anesthetized two-photon experiment using the same procedures as described previously [58]. Briefly, during the experiment ferrets were induced with 40 mg/kg ketamine and 0.05 mg/kg atropine IM and maintained on isoflurane anesthesia, and paralyzed using pancuronium bromide (0.15 mg/kg/hr). Continuous monitoring of a range of vital parameters (heart rate, SpO2, EKG, EtCO2 and the EEG) ensured adequate anesthetic depth during the experiment. A custom-made stainless steel imaging chamber was cemented to the skull centered on the virus injection site. The bone and dura over the virus injection site were then removed, the brain covered with a thin layer of agarose (type III, Sigma-Aldrich) and a coverslip. Imaging was performed at 920nm using a Coherent Ti:Sapphire laser coupled to a two-photon microscope from Neurolabware.

#### 5.4.8. Porcine eye experiments

The study was conducted with approval by the Administrative Panel for Laboratory Animal Care at Stanford University and in accordance with the Guide for the Care and Use of Laboratory Animals at an AAALAC-accredited facility. A 6-month-old female Yucatan minipig was obtained from an approved vendor (Premier BioSource, CA), acclimated for at least 3 days, and group housed under standard conditions.

Before anesthesia, the animal was fasted for 12-18 hours. The animal was sedated with ketamine (5 mg/kg), dexmedetomidine (0.03 mg/kg), and butorphanol (0.2 mg/kg) intramuscularly (IM) and maintained with mask isoflurane (1-4%) in 100% oxygen. Heart rate, respiratory rate, blood oxygen saturation, body temperature, and jaw tone were monitored continuously and recorded every 15 minutes.

The right eye was topically treated with tetracaine hydrochloride ophthalmic solution (0.5%). The right orbital area was clipped and aseptically prepared with saline and ophthalmic betadine (5%). Using sterile technique, a 30-gauge insulin syringe was used to aspirate 150 µL of aqueous fluid. With the guidance of Castroviejo calipers, AAV2-CAG-GFP (50 µL, 1.5 x 10^13^ GC/mL, Addgene 37825-AAV2) was injected intravitreally 4 mm posterior to the temporal aspect of the limbus, and AAV2-CAG-TdTomato (100 µL, 5.3 x 10^12^ GC/ml, Addgene 59462-AAV2) was injected intravitreally 4 mm posterior to the nasal aspect of the limbus. Neomycin, polymyxin B sulfate, and bacitracin zinc ophthalmic ointment was topically applied to the eye. Anesthesia was reversed with atipamezole (0.35 mg/kg) IM, and the animal was recovered.

Seven weeks later, the animal was sedated with Telazol (6 mg/kg) IM and euthanized with pentobarbital sodium and phenytoin (8 mL; 390 mg/mL, 50 mg/ml) IV. Subsequently, the right eye was harvested for two-photon imaging.

Two-photon imaging was performed using the 20 mm working distance objective on an Ultima IV two-photon microscope (Bruker) equipped with 2 GaAsP PMTs and a custom fiber nosepiece. Laser excitation consisted of a variable wavelength laser set to 920 nm, and a steady state laser at 1040 nm (Spectra-Physics). The enucleated eye was placed under the objective with hydroxypropyl methylcellulose gel (Ocular Vision) and a 1.5 coverslip on top.

## Acknowledgments

We thank Special Optics for manufacturing the objective. We thank George Chen for discussing the optical design and Antonio Perez for the mechanical design at Special Optics. We thank collaborators Stephen Macknik, Tom Franken, Stephen Van Hooser, Joe Culver, and Ben Huang for early support of the project and valuable input.

## Contributions

KN motivated the project, and worked with CHY and SLS to develop the specifications. CHY designed the objective, coordinated fabrication, and characterized it. YY and CHY performed mouse imaging. ITS and LA, along with CHY and YY, performed the dendritic spine and bouton imaging. JTC, JB, and DF performed the ferret and tree shrew imaging. AHM, HD, JAC, and MH performed the dual prism-widefield imaging in mice. JK and AR performed the cochlea imaging. AV designed the imaging system used in the marmoset imaging. DGCH and SOC performed marmoset imaging experiments. RV, BRN, and KN performed the ferret experiments with objective comparisons. BKY and KH performed the porcine eye imaging. DMF, AL, RP, and PK performed the three-photon imaging. GBS provided feedback on mechanical design, usability, and performance. CHY and SLS wrote the paper with contributions from all authors. SLS led the project.

## Data Availability Statement

The design of the objective is fully open source and full specification are detailed in this report. Data underlying the results from imaging in animals are available from the authors upon request.

## Code availability

The code used in this work is already publicly available or detailed in the methods section. If additional materials are required for replication, the authors invite such requests.

## Supplementary Videos

Video 1: Z-stack of *in vivo* calcium imaging.

The z-plane range spans from the brain surface to the depth of 500 *µ*m. Frame size = 1024 x 1024 pixels. Imaging power = 80 mW.

Video 2: *In vivo* calcium imaging over a 1.7 x 1.7 mm FOV.

Frame rate = 15.4 frames/s. Frame size = 1536 x 1536 pixels. Imaging power = 60 mW. Imaging depth = 250 *µ*m.

Video 3: *In vivo* three-photon imaging z stack from the mouse visual cortex and white matter below using the Cousa objective lens. The XY field-of-view shown represents a region of 0.7 mm × 0.7 mm. The depth range spans 0.270 mm to 0.945 mm from the pial surface and thus starts in cortical layer 2/3 and ends below the white matter. The lumen of blood vessels is visible (shown in magenta) after a retro-orbital injection of Texas Red Dextran. Apical dendrites (distinct puncta) and white matter axonal fibers are visible with THG (shown as cyan).

## 6. Supplementary Material

**Fig. S1.**
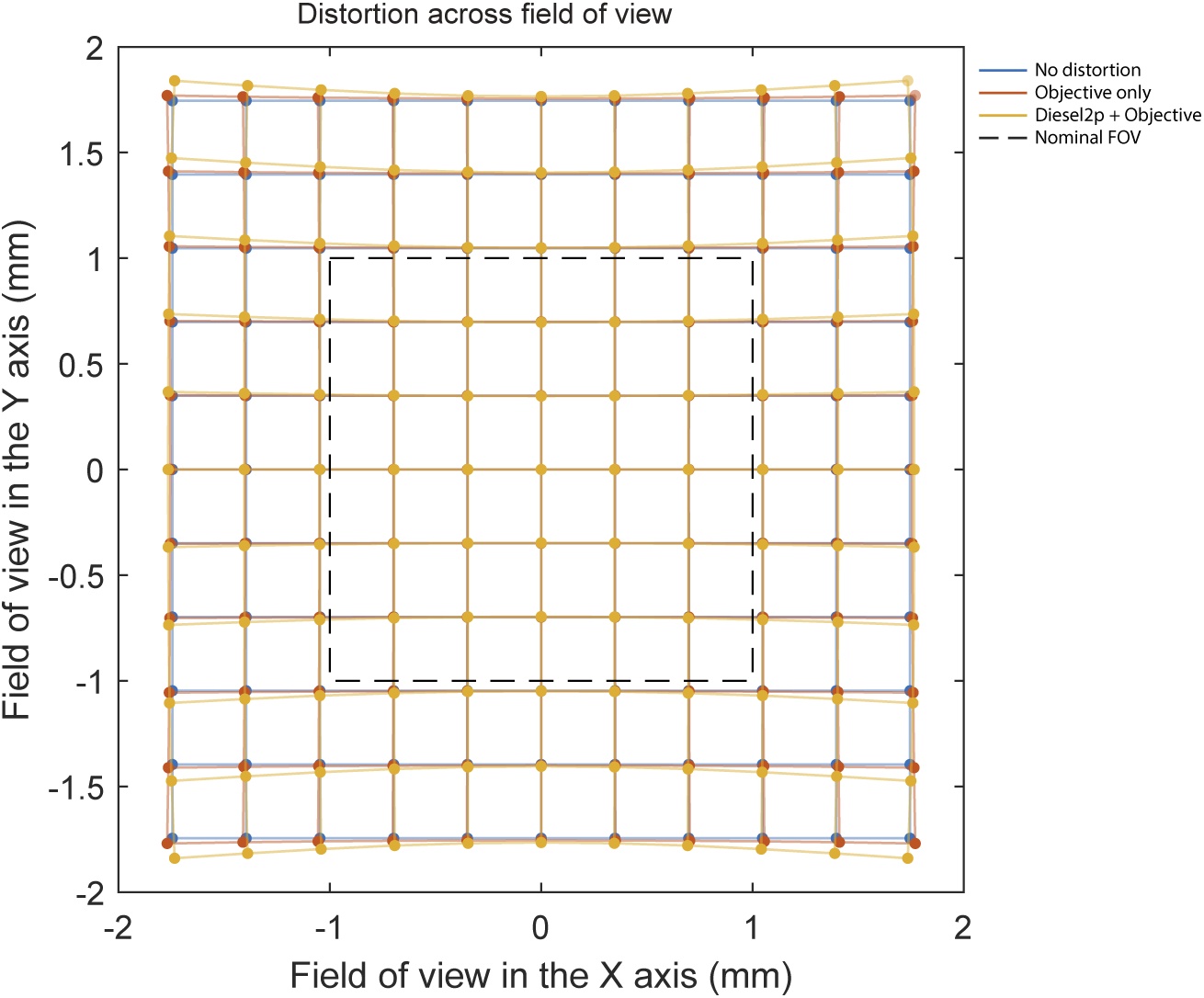
The diagram shows the distortion of the field of view simulated from the Zemax model. The blue grid shows the undistorted (square) FOV for comparison. The orange grid shows the shape of the FOV for the Cousa objective, showing a pincushion distortion. The yellow grid shows the shape of the FOV formed by the sum of the Diesel2p scan engine and the Cousa objective, showing a combination of the pincushion and barrel distortion. The shape of the simulated distortion (in yellow) matches with the distortion of the experimental measurement shown in Fig. 1c. Note that there is nearly zero field distortion in the 2 x 2 mm^2^ nominal FOV (dashed black square).

**Fig. S2.**
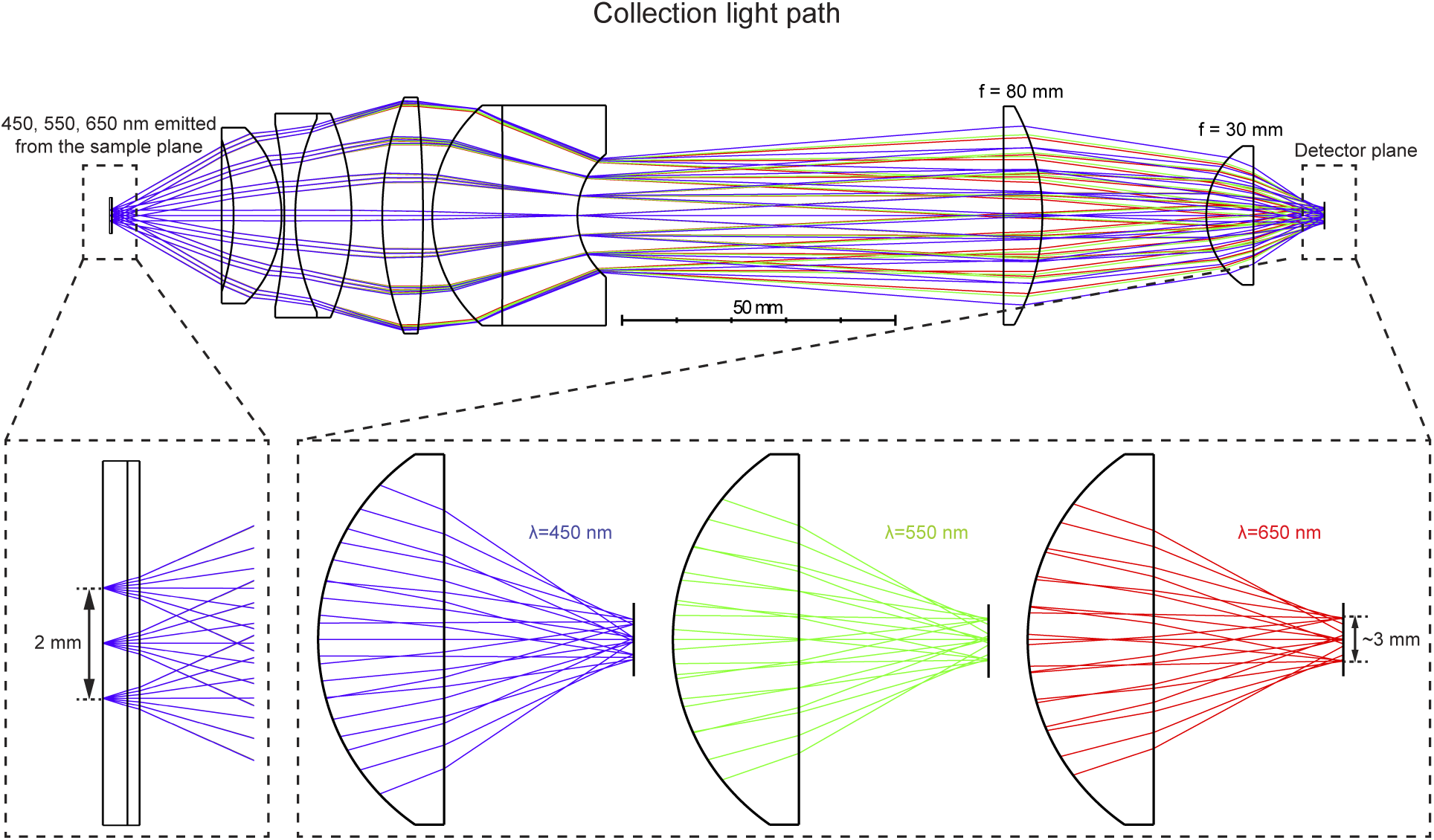
A schematic shows the ray-traces in the collection light path for the Cousa objective. Fluorescence photons with wavelengths in the visible band (450 nm, 550 nm, and 650 nm) are emitted from the imaging plane within the 2 mm nominal FOV (zoom in view on the left), passing through the Cousa objective. These photons are further collected and guided by the optical relay, and reach at the cathode of the photomultiplier (PMT). The optical relay is constructed simply with two off-the-shelf singlet lenses from OptoSigma (011-2570-a55) and Thorlabs (LA1805-A). The zoom in view in the bottom-right shows that all of the emitted photons (rays) in the visible spectrum reach at the detector surface within a 3 mm wide spread.

**Fig. S3.**
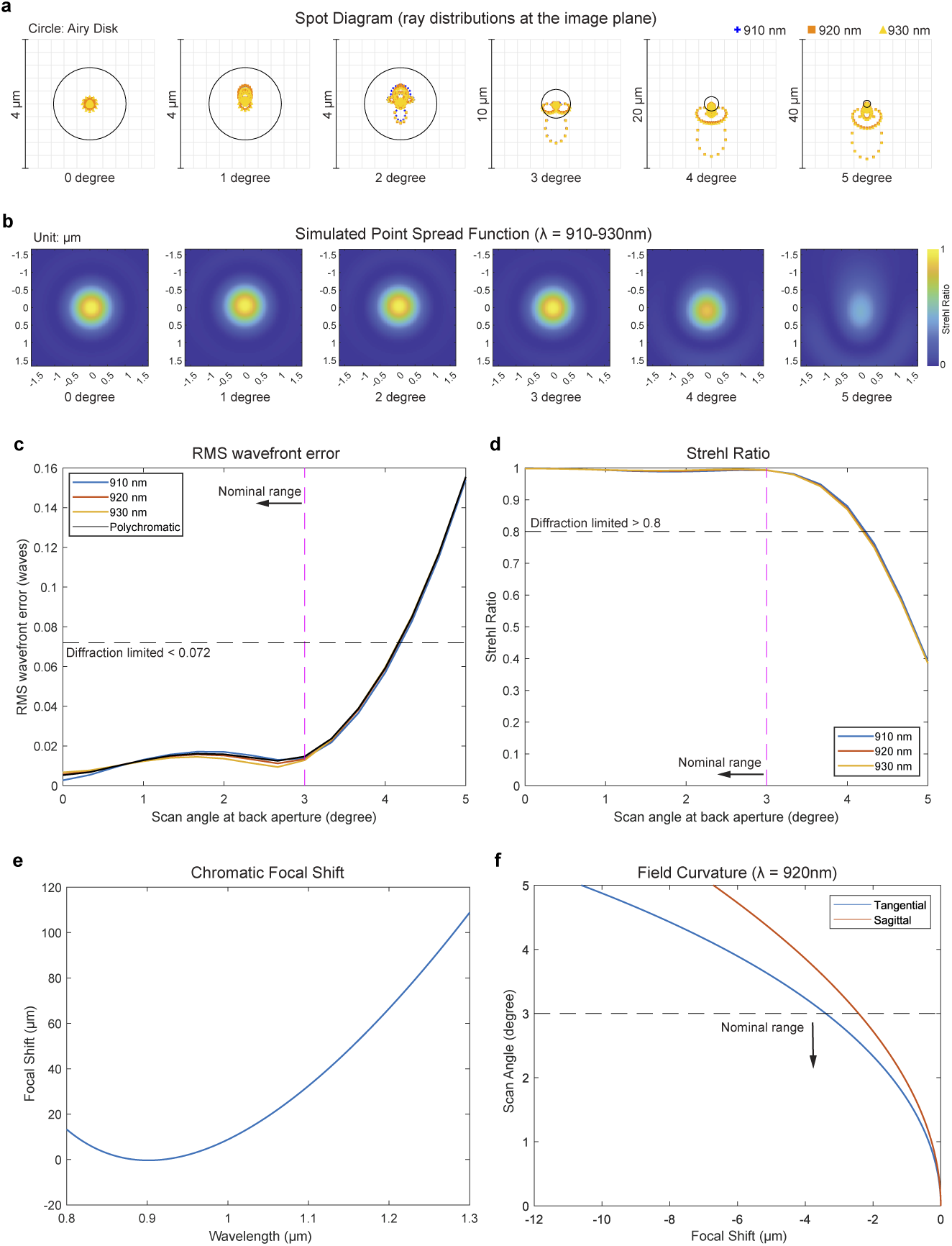
Additional information on nominal performance. (a) The spot diagram shows the distribution of rays at the imaging plane coming from a scan angle of 0^◦^, 1^◦^, 2^◦^,3 ^◦^, 4^◦^, 5^◦^, aiming at the objective’s back aperture with a beam diameter of 20 mm, and with wavelengths of 910 nm, 920 nm, and 930 nm. (b) The simulated point spread functions from Zemax at the 5 scan angles are shown. The root-mean-square wavefront error (c) and the Strehl ration (d) as a function the scan scan angle at the back aperture is shown for the wavelength of 910nm, 920nm, 930nm, and all three combined (polychromatic). The horizontal dashed line indicates that the curve below 0.072 in the RMS wavefront error plot and above 0.8 in the Strehl plot is diffraction limited. The vertical dashed line in (c) and (d) shows the nominal range of scan angle is smaller than 3 degrees. (e) The dependence of focal shift on the wavelength in the center of FOV is shown. (f) The depth of the tangential and sagittal focus over a scan angle of 5 degrees at 920 nm is shown. The horizontal dashed line shows the nominal range of the scan angle < 3^◦^. There is only 4 *µ*m depth difference in the center (0^◦^) and in the edge (3^◦^) of the FOV within the nominal scan range.

**Fig. S4.**
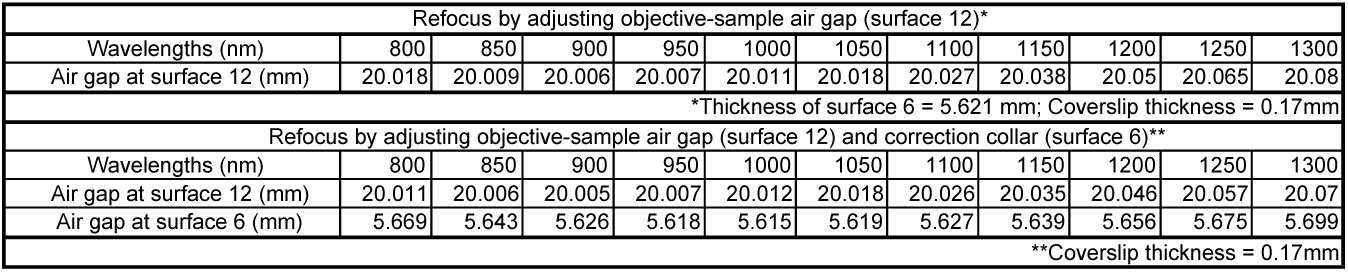
Thickness of air gaps for optimal performance at different wavelengths. The nominal thicknesses of air gaps used for different wavelengths. The upper portion of table lists the distance of the objective-to-sample gap (Surface 12) when this single air gap is adjusted for different wavelengths. The lower part of the table shows the thicknesses of the objective-to-sample gap (Surface 12) and the correction collar gap (Surface 6) when both of these two air gaps are adjusted.

**Fig. S5.**
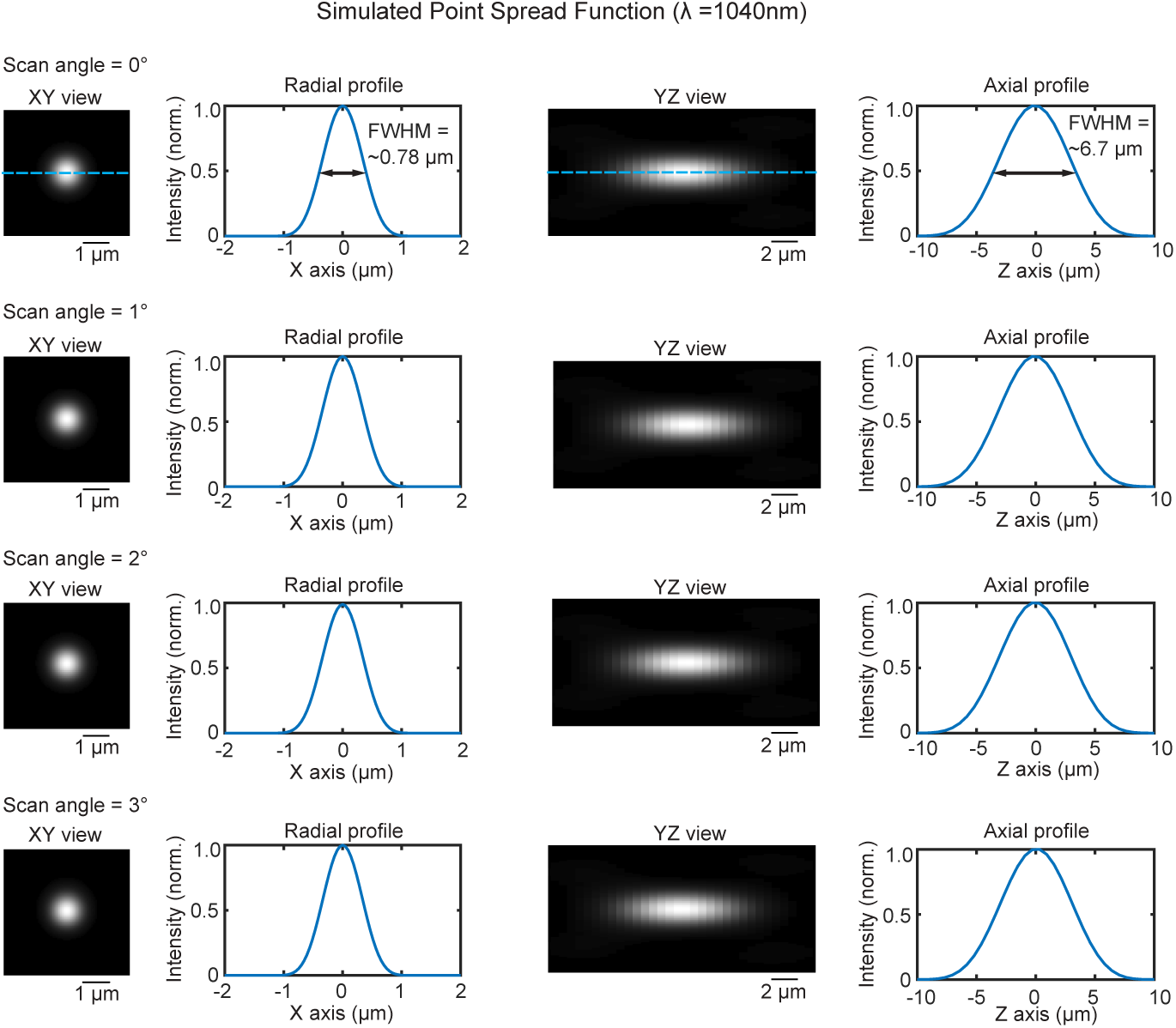
Focusing 1040 nm light. Across scan angles from 0 to 3 degrees, the focus of 1040 nm light is < 0.8 *µ*m laterally and < 7.0 *µ*m axially.

**Fig. S6.**
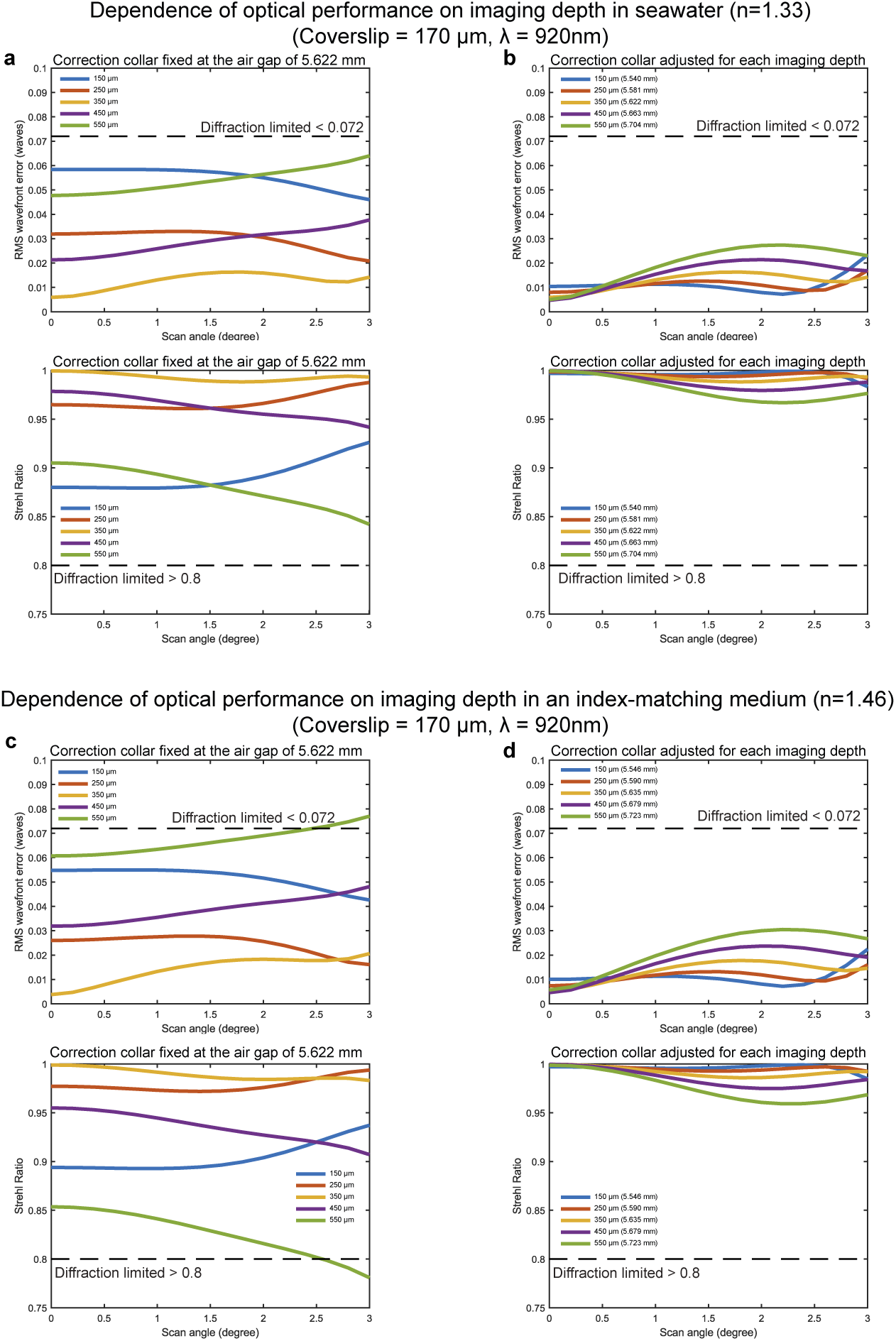
Dependence of performance on the imaging depth and refractive index of the sample medium as well as the compensation using the correction collar. The root-mean-squared (RMS) wavefront error (top) and the Strehl ratio (bottom) as a function of the scan angle at the back aperture at different imaging depths inside seawater (n=1.33) are shown with the correction collar fixed at a air gap of 5.622 mm in (a), and with the correction collar optimized for different imaging depths in (b). Different imaging depths ranging from 150-550 *µ*m are color coded. The same plots inside an index-matching medium (n=1.46) are shown as (a) with the correction collar fixed at a air gap of 5.622 mm in (c), and with the correction collar optimized for different imaging depths in (d).

**Fig. S7.**
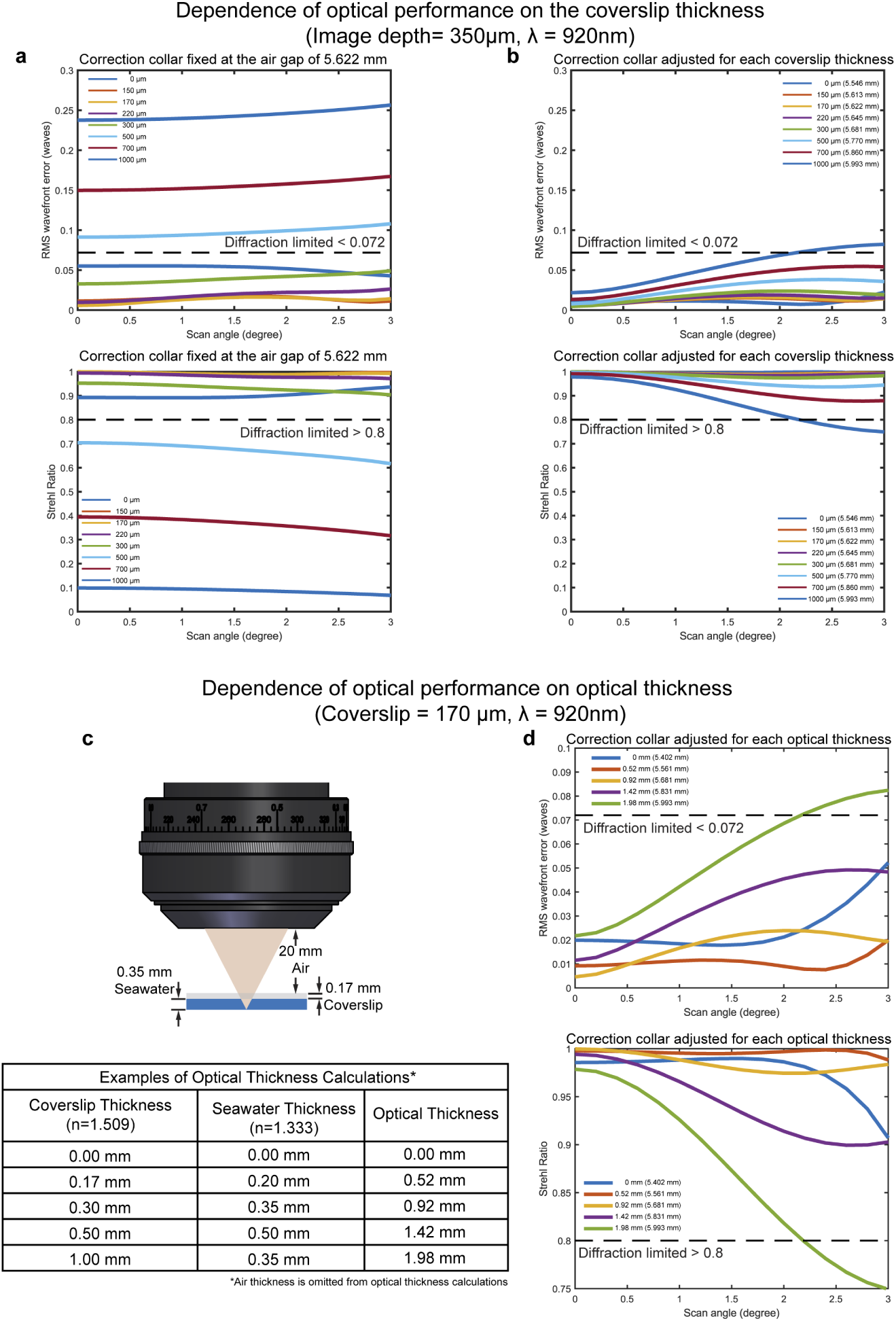
Dependence of performance on coverslips of different thickness and the compensation using the correction collar. The root-mean-squared (RMS) wavefront error (top) and the Strehl ratio (bottom) as a function of the scan angle at the back aperture and coverslips of different thickness are shown with the correction collar fixed at a air gap of 5.622 mm in (a), and with the correction collar optimized for coverslips of different thickness in (b). The thickness of coverslips ranging from 0-1000 *µ*m are color coded. (c) The schematic shows the Cousa objective focuses light through a 0.17 mm thick coverslip and 0.35 mm of seawater. The optical thickness of the non-air elements between the objective and the focal plane can be calculated: multiply the thickness of a material (ignoring air) in millimeters by its refractive index, and sum over all of these products. For example, the optical thickness shown in the schematic is 0.723 (0.17 mm x 1.509 + 0.35 mm x 1.333). The table below shows combinations of the coverslip thickness and the seawater thickness, and the resultant optical thickness for each combination. The correction collar can be adjusted to compensate for the optical thickness over the range of 0 - 2mm, and maintain the diffraction-limited performance. (d) The root-mean-squared (RMS) wavefront error (top) and the Strehl ratio (bottom) as a function of the scan angle at the back aperture are shown when the correction collar is optimized for different optical thickness of non-air materials. Each curve is color coded for different optical thickness shown as the number before the parenthesis. The number in the parenthesis is the optimal thickness of air gap to achieve the best optical performance.

**Fig. S8.**
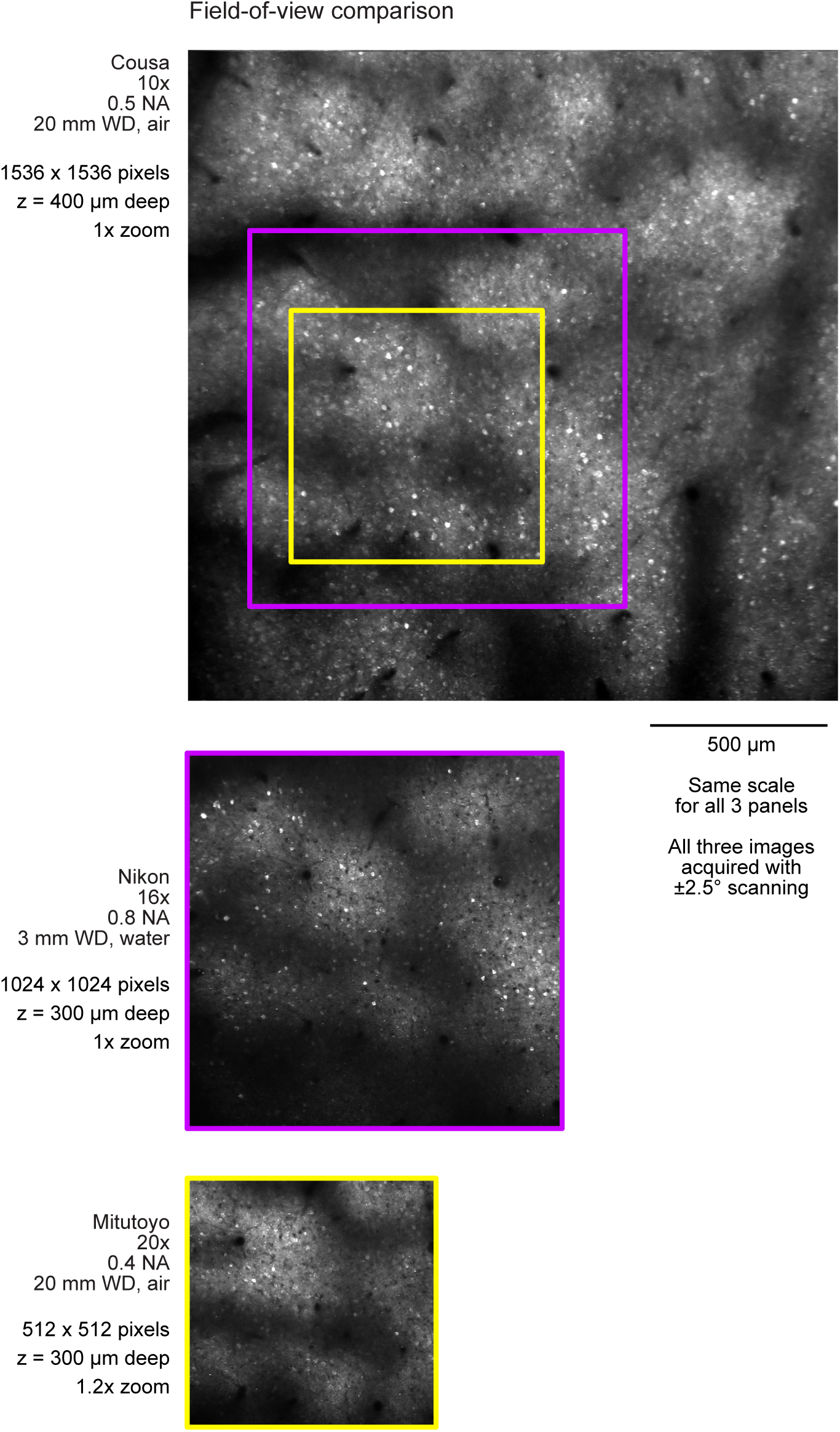
Comparing the field-of-view (FOV) between the Cousa and two conventional objectives.

**Fig. S9.**
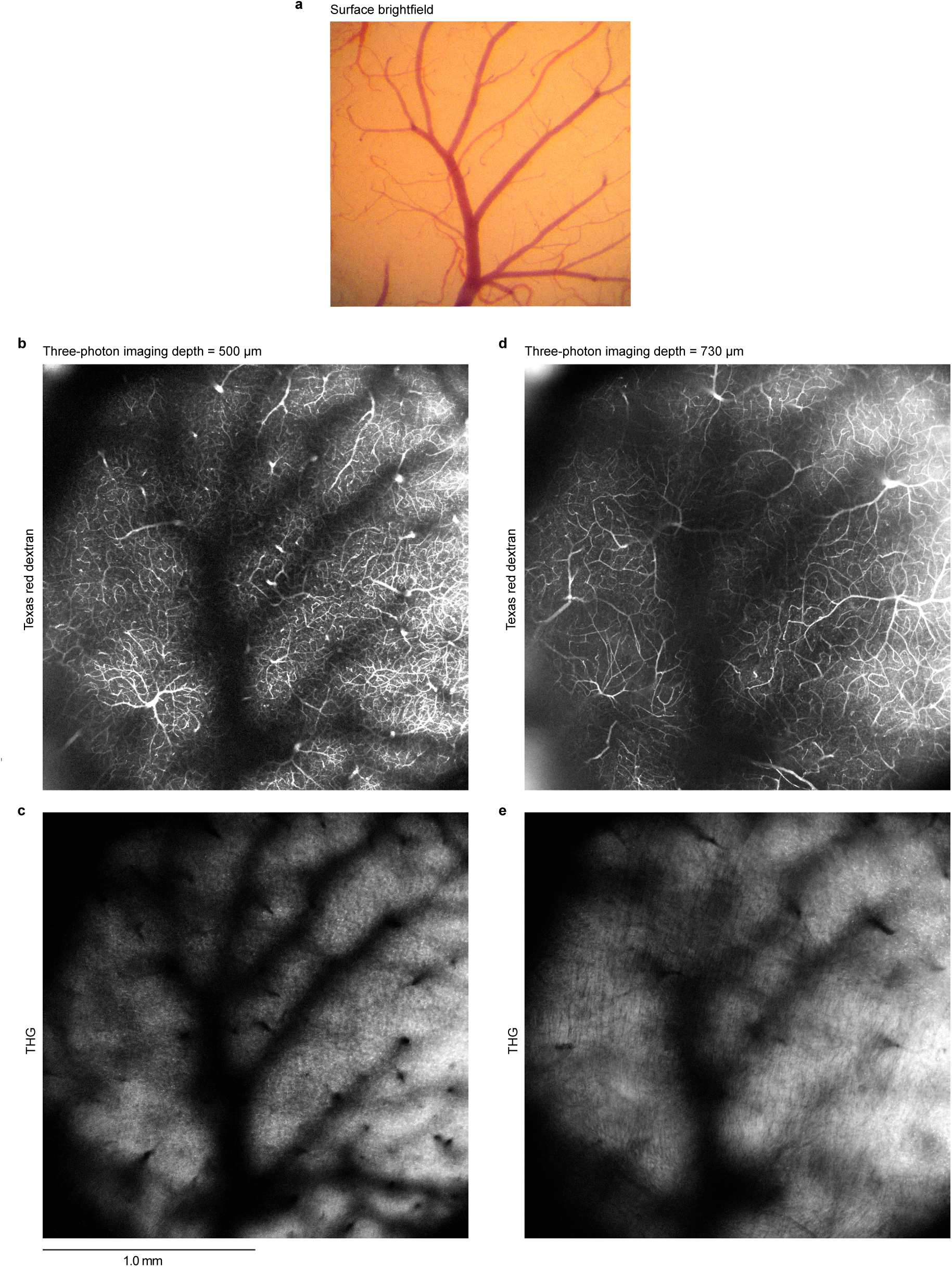
Large field-of-view three-photon imaging with the Cousa objective. (a) Brightfield image of a craniotomy over the mouse visual cortex, viewed through a surgical dissecting microscope. No dyes were present in the circulation at this phase of the experiment. Arteries and veins can be distinguished based on small difference in hue—arteries shown in orange and veins in purple. (b,c) Single z plane three-photon images (dual channel recording) from mouse visual cortex, 500 µm below the pial surface showing blood vessels labeled with Texas Red dextran (panel b) and THG signals (panel c). (d, e) Single z plane three-photon images (dual channel recording) from mouse visual cortex, 730 µm below the pial surface showing blood vessels labeled with Texas Red dextran (panel d) and THG signals (panel e).

